# Magainin 2 and PGLa in Bacterial Membrane Mimics I: Peptide-Peptide and Lipid-Peptide Interactions

**DOI:** 10.1101/664359

**Authors:** Michael Pachler, Ivo Kabelka, Marie-Sousai Appavou, Karl Lohner, Robert Vácha, Georg Pabst

## Abstract

We addressed the onset of synergistic activity of the two well-studied antimicrobial peptides magainin 2 (MG2a) and PGLa using lipid-only mimics of Gram-negative cytoplasmic membranes. Specifically, we coupled a joint analysis of small-angle X-ray and neutron scattering experiments on fully hydrated lipid vesicles in the presence of MG2a and L18W-PGLa to all-atom and coarse-grained molecular dynamics simulations. In agreement with previous studies both peptides, as well as their equimolar mixture, were found to remain in a surface-aligned topology upon membrane insertion and to induce significant membrane perturbation as evidenced by membrane thinning and hydrocarbon order parameter changes in the vicinity of the inserted peptide. These effects were particularly pronounced for the so called synergistic mixture of 1:1 (mol/mol) L18W-PGLa/MG2a and cannot be accounted for by a linear combination of the membrane perturbations of two peptides individually. Our data are consistent with parallel heterodimers forming at much lower concentrations than previously considered, but which do not induce a synergistic leakage of dyes. Our simulations further show that the heterodimers interact via salt bridges and hydrophobic forces, which apparently makes them more stable than putatively formed antiparallel L18W-PGLa and MG2a homodimers. Moreover, dimerization of L18W-PGLa and MG2a leads to a relocation of the peptides within the lipid headgroup regime as compared to the individual peptides. The early onset of dimerization of L18W-PGLa and MG2a at low peptide concentrations consequently appears to be key to their synergistic dye-releasing activity from lipid vesicles at high concentrations.

**STATEMENT OF SIGNIFICANCE:** We demonstrate that specific interactions of the antimicrobial peptides MG2a and PGLa with each other in POPE/POPG bilayers lead to the formation of surface-aligned parallel dimers, which provide already at low peptide concentrations the nucleus for the peptides’ well-known synergistic activity.

## INTRODUCTION

The steady increase of antibiotic resistance of pathogenic bacteria combined with the decline of approved antimicrobial agents is considered to be a severe threat to global health. In view of these developments, considerable research efforts have been devoted to understanding the mode of action of antimicrobial peptides (AMPs), considered as alternative for the development of novel antibiotics. AMPs are effector molecules of the innate immune system, whose main targets are bacterial membranes. AMPs kill bacteria within minutes, which makes it more difficult for bacteria to develop resistance mechanisms (for review see e.g. (1, 2)). Applying diverse biophysical techniques on lipid-only membrane mimics, several interaction models have been conceived for AMPs (1, 3, 4). Pore-formation is arguably the most widely discussed membrane disruptive mechanism.

Choosing a specific membrane mimic requires a delicate balance between experimental or computational tractability and physiological relevance. This leads to a search for the minimum realistic lipid mixture which yields a similar response to AMPs as in live bacteria. Phosphatidylethanolamine, phosphatidylglycerol, and cardiolipin represent the main lipid components of Gram-negative cytoplasmic membranes. Yet, we demonstrated previously that lipid bilayers composed of palmitoyl-oleoyl-phosphatidylethanolamine (POPE) and palmitoyl-oleoyl-phosphatidylglycerol (POPG) (molar ratio: 3:1) respond similarly to PGLa and Magainin 2-amide (MG2a), in agreement with their antimicrobial activity toward *Escherichia coli* K12 (5), despite the lack of cardiolipin or other cell wall components. Hence POPE/POPG (3:1 mol/mol) bilayers appear to be valid first order mimics for biophysical studies on the activity of AMPs with Gram-negative bacteria.

Studying the activity of PGLa and MG2a is of specific interest due to synergistic effects, first described by Matsuzaki and coworkers (6–8). Equimolar mixtures of the two peptides reduced the minimum inhibitory concentration of the individual peptides in *E. coli* K12 by about one order of magnitude. The molecular mechanism of the observed synergy remains controversial, however. Originally, synergism was associated with the formation of a transmembrane pore with a 1:1 peptide stoichiometry (6–8). In contrast, solid state NMR experiments from the Ulrich and Bechinger groups demonstrated that MG2a never adopts a transmembrane topology and that PGLa may insert perpendicularly in the presence of MG2a into the membrane only in phosphatidylcholine-enriched or short chain disaturated phosphatidylethanolamine-enriched bilayers, respectively (9–13). None of these latter lipids are of significance for Gram-negative cytoplasmic membranes, however.

The dependence of MG2a mediated PGLa insertion into bilayers was attributed to the intrinsic lipid curvature, which leads to a tight packing of the bilayer’s polar-apolar interface in the case of cone-shaped lipids, such as POPE, and increases the energy barrier for peptide translocation (5, 14). Of recent, this was refined by considering also details of hydrocarbon chain configurations (13).

The topologies of PGLa and MG2a within the synergistic regime as proposed from the above mentioned data differ significantly. Zerweck et al. postulated a pore formed by a tetrameric heterocomplex of transmembrane PGLa and surface aligned MG2a, which is stabilized by intimate Gly-Gly contacts between antiparallel PGLa dimers and C-terminal interactions between PGLa and MG2a (12). This contrasts, however, the surface topology of both peptides reported in POPE enriched bilayers (11, 13). Notably some of the observed effects on peptide topology might be also related to the relatively low water content of solid state NMR experiments. We are thus currently lacking insight that would explain how PGLa and MG2a remain surface bound, but disrupt membrane at the same time synergistically (5).

We therefore performed a comprehensive study using a broad selection of experimental and computational tools in order to reveal the effects of PGLa and MG2a in fully hydrated POPE/POPG (3:1 mol/mol) bilayers on nanoscopic to macroscopic length scales. Specific care was given to ensure that conditions allow a direct comparison to our previously reported leakage experiments, including the use of L18W-PGLa instead of native PGLa (5). Note that L18W-PGLa was reported to behave analogously to native PGLa (6).

Due to the large number of produced data, we decided to present the results in a paper series. The present paper focuses on low peptide concentrations, i.e. where L18W-PGLa/MG2a mixtures do not cause synergistic dye-release from POPE/POPG vesicles (5). This allowed us to investigate the peptides influence on the membrane structure in great detail by using combined joint small-angle X-ray and neutron scattering (SAXS/SANS) as well as all-atom and coarse-grained molecular dynamics (MD) simulations. We found, for example, that membrane-mediated interactions between the two peptides lead to an early onset of dimerization causing a shift of L18W-PGLa from slightly below to slightly above the lipid’s glycerol backbone, while remaining surface aligned. This effect leads to a perturbation of membrane structure, which is more pronounced than in the case of non-interacting individual peptides. The resulting remodelling of membrane structure thus appears as a precursor to synergistic dye release at higher peptide concentrations.

## MATERIALS AND METHODS

### Lipids, Peptides and Chemicals

POPE and POPG were purchased from Avanti Polar Lipids (Alabaster, AL, USA, purity>99%) as powder and used without further purification. L18W-PGLa (GMASKAGAIAGKIAKVAWKAL-NH_2_) and MG2a (GIGKFLHSAKKFGKAFVGEIMNS-NH_2_) were obtained in lyophilized form (purity >95%) from PolyPeptide Laboratories (San Diego, CA, USA). Deuterium dioxide (purity 99.8 atom %) and HEPES (purity >99.5) were purchased from Carl Roth (Karlsruhe, Baden-Wüttenberg, Germany). All other chemicals were obtained from Sigma-Aldrich (Vienna, Austria) in *pro analysis* quality. Lipid stock solutions for sample preparation were prepared in organic solvent chloroform/methanol (9:1; v/v); lipid concentration was determined using a phosphate assay (15). Peptide stock solutions were prepared in 10 mM HEPES 140 mM NaCl buffer solution (pH 7.4).

### Sample Preparation

Lipid thin films were prepared by mixing appropriate amounts of lipid stock solutions to obtain samples composed of POPE:POPG (3:1, mol/mol), followed by solvent evaporation under a nitrogen stream at 35°C and overnight storage in a vacuum chamber. Dry lipid films were hydrated in 10 mM HEPES, containing 140 mM NaCl (pH 7.4). For neutron experiments the H_2_O/D_2_O ratio in the buffer was varied as detailed below. Hydrated samples were equilibrated for one hour at 55°C followed by 8 freeze-and-thaw cycles using liquid N_2_ and intermittent vortex mixing. Large unilamellar vesicles (LUVs) were obtained by 31 extrusions with a hand held mini extruder (Avanti Polar Lipids, Alabaster, AL) using a 100 nm pore diameter polycarbonate filter. Vesicle size and polydispersity was determined via dynamic light scattering using a Zetasizer NANO ZSP (Malvern Instruments, Malvern, United Kingdom). LUVs were again phosphate assayed and mixed with appropriate amounts of peptide stock solution to obtain peptide/lipid (P/L) molar ratios in the range of P/L = 1/400 – 1/50. LUVs were equilibrated at a given peptide concentration for 4 days prior to measurement.

### Small-angle X-ray scattering

Small-angle X-ray scattering (SAXS) data were collected at the SWING beamline (Soleil, Saint-Aubin, France) using X-ray photons of wavelength *λ* = 10 Å and an Eiger 4M detector (Dectris, Baden-Daetwill, Switzerland). Samples were manually loaded in 1.5 mm path length quartz capillaries and mounted in a capillary holder, whose temperature was controlled with a circulating water bath. The sample-to-detector distance (SDD) was set to 1 m, which allowed us to cover scattering vectors in the range from *q* = 0.0098 Å^−1^ to 0.9 Å^−1^. Data correction (integration, normalization and background subtraction) was performed using the software Foxtrot (Xenocs, Sassenage, France).

### Small-angle neutron scattering

Neutron scattering experiments (SANS) were performed at KWS-1 (FRM II, Munich-Garching, Germany (16). Using 2D scintillation detector, a wavelength of 5 Å (Δ*λ* /*λ* = 0.1) and SDD = 1.21 m and 7.71 m, allowed us to cover a *q*-range from 0.005 − 0.42 Å^−1^. Samples were kept in 1 mm path length quartz cuvettes QX-404 (Helmma, Jena, Germany) and equilibrated at 35°C, using a circulating water bath. Used contrast conditions were 100, 75 and 50 % v/v D2O/H2O. Data correction was performed using the QTIKWS software from JCNS (Gaching, Germany).

### Joint SAXS/SANS analysis

For the P/L ratios used in the present study no peptide-mediated aggregation of LUVs was observed, i.e. scattering data did not exhibit any Bragg peak scattering. This allowed us to treat SAXS/SANS data in terms of dilute particle scattering, that is we were able to neglect any contribution from LUV-LUV or bilayer-bilayer positional correlations. Moreover, we focused our analysis on peptide-induced changes of the membrane structure by analyzing scattering data for *q* > 0.05 Å^−1^. In this *q*-range overall vesicle size and morphology do not contribute (17), which allowed us to model the scattered intensities as *I*(*q*) ∝ |*F*_FB_(*q*)|^2^/*q*^2^, where *F*_FB_(*q*) is the flat bilayer form factor.

The form factor was derived within the framework of the scattering density profile (SDP) model (18). In brief, the SDP analysis is based on a composition-dependent parsing of membrane structure into quasi-molecular fragments, whose distribution along the bilayer normal is described in terms of Gaussian-type volume probability functions. Parsing is guided by MD simulations (for a recent review, see e.g., (19)). The volume probability functions can be easily scaled with the neutron or X-ray scattering lengths of each group entailing a joint SAXS/SANS analysis that combines the differently contrasted samples into one underlying membrane structure. Analogous strategies have been reported previously, see e.g. (20–23).

#### Parsing

Considering previously reported SDP data for POPE (24), POPG (25), and by using all-atom MD simulations (see below) the lipid part of the membrane structure was parsed into the ethanolamine (ENX), glycerol (PG2), phosphate (PO_4_), carbonyl glycerol (CG), CH, CH_2_, and CH_3_ groups (see Fig. 1 and supplementary Fig. S1). Following our previous SDP analysis of coexisting lipid domains (26, 27), we combined the individual groups of POPE and POPG into one hybrid lipid structure, using molecular averaging. In particular, we paired all hydrocarbon, CG and PO_4_ groups, assuming that they align at the same transbilayer position. This is reasonable considering the identical hydrocarbon chain composition of both lipids. That is, only the ENX and PG2 groups were adjusted independently.

**Figure 1:**
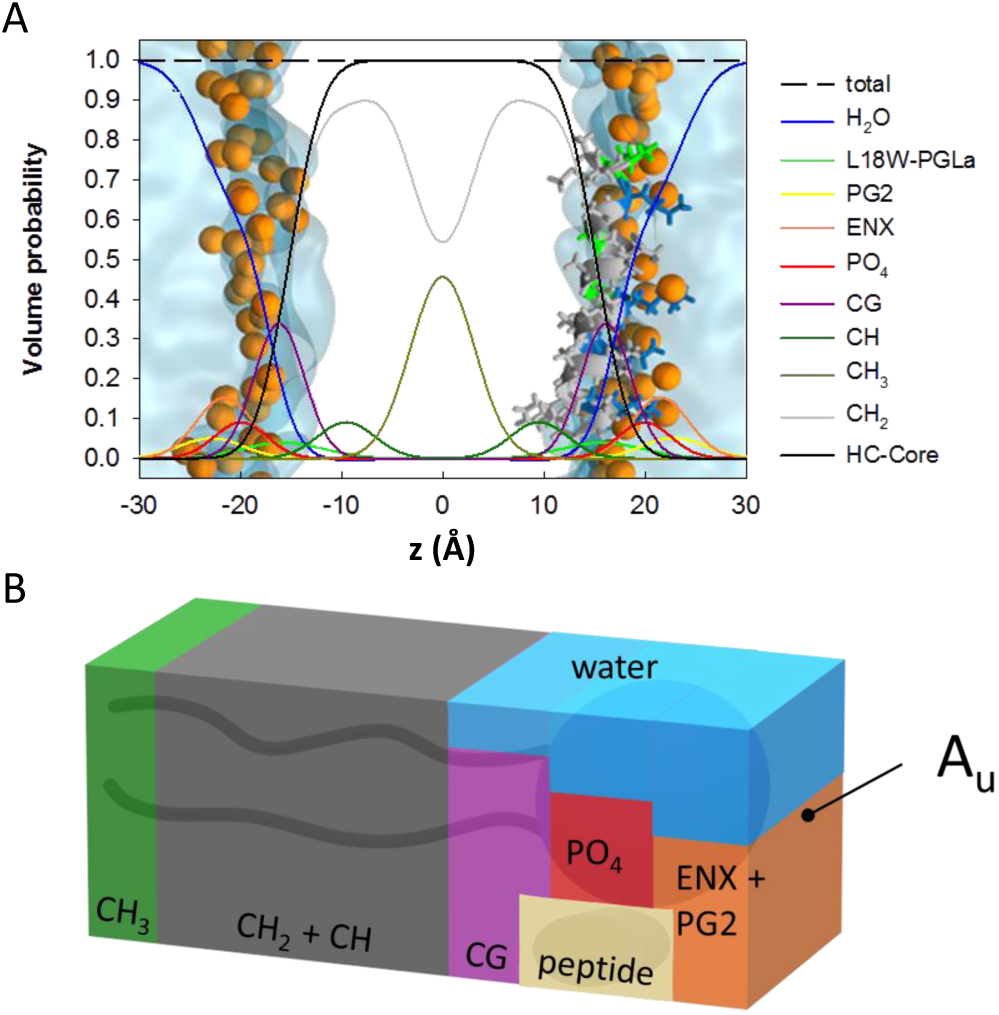
Schematic of the applied SDP model. Panel A shows the distribution functions of the different quasimolecular groups (see also Fig. S1). The spheres in the overlayed MD simulations represent phosphor atoms. Panel B gives a schematic of the unit cell of cross sectional area *A*_U_ with different contributions from lipid, peptide and water.

Based on experimental evidence for surface-aligned topologies of MG2a and L18W-PGLa in POPE/POPG (3:1 mol/mol) (13), as well as our own MD simulations results, we modelled the contribution of the peptides by a single Gaussian volume probability function centered at *z*_P_ in the headgroup regime. Further, we assumed that peptides (i) distribute equally in both leaflets and (ii) fully partition into the lipid membrane. The first assumption is motivated by the long sample equilibration times, during which peptides are able to translocate spontaneously through the bilayer. The second assumption is corroborated by the absence of scattering from unbound peptides in our data (28) and previous partitioning experiments (29). For the equimolar mixture, we combined L18W-PGLa and MG2a into a single Gaussian as supported by MD simulations (see below).

The individual volume distribution functions are detailed in the SI. The area per unit cell *A*_U_ is a common scaling factor for all distribution functions and therefore was chosen as fit parameter (see also (27)). From the analysis we determined several structural parameters, such as, e.g., the Luzzati bilayer thickness (30)

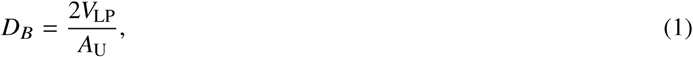

where *V*_VP_ is the total volume of the lipid/peptide unit cell (Eq. S9), the hydrocarbon chain length 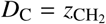, where 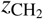 is the outer terminal position of the methylene group, and the distance from the CG to PO_4_ groups 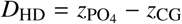.

#### Data fitting

To obtain robust fitting parameters and estimates for their uncertainties every data set was fitted 400 times using a genetic algorithm (31) with random starting parameters. This gave us Gaussian-like distributions for each adjustable parameter. We report the average (center of mass) of these distributions; uncertainties where calculated from second moments. For details regarding constraints and cost function, see the SI.

### Molecular dynamics simulations

Molecular dynamics (MD) simulations were performed using GROMACS version 2016.2 (32, 33).

#### All-atom simulations

##### Simulation Settings

Protein and solvent molecules were described by Amberff99SB-ILDN (34, 35), and lipids by Slipids (36, 37) force fields. The simulation time step was set to 2 fs. A temperature of 308.15 K was maintained by using a Nosé-Hoover thermostat (38–40), with a coupling constant of 0.5 ps. Two separate coupling groups for protein-lipid and solvent atoms were used. 3D periodic boundary conditions were applied and a Parrinello-Rahman barostat (41, 42) with semi-isotropic coupling scheme was employed for keeping the pressure at 1 bar with a coupling constant of 2 ps. Long-ranged electrostatic interactions were treated with particle mesh Ewald method (43) with the real-space cutoff set to 1.2 nm.Lennard-Jones interactions were cutoff at 1.2 nm. All bonds were constrained using the LINCS algorithm; long-range dispersion corrections (44) were applied for energy and pressure.

##### System Preparation

Lipid bilayers composed of 192 POPE and 64 POPG molecules were assembled in the *XY* -plane by distributing lipids equally in both leaflets using the CHARMM-GUI interface (45). The system was hydrated by more than 40 water molecules per lipid and NaCl ions were added at 130 mM concentration. The initial box dimensions were 8.9 × 8.9 × 8.5 nm. MG2a and L18W-PGLa were prepared in *α*-helical conformation.

The following starting configurations were considered. System (1): A single peptide was placed into each membrane leaflet in a surface-aligned topology. System (2): A parallel L18W-PGLa/MG2a heterodimer was placed into each membrane leaflet in a surface-aligned topology.

All systems were equilibrated using similar protocols. Firstly, energy minimization was performed using the steepest descent algorithm. Then, an equilibration with positional restraints on peptide backbone was performed for 60 ns followed by an equilibration with dihedral restraints, in order to maintain the peptide’s secondary structure. The length of simulations with dihedral restrains were 105 ns (system (1)), or 180 ns (system (2)). Finally, unrestrained production dynamics simulation were performed for 500 ns.

### Coarse-grained Simulations

#### Simulation Settings

Computationally-efficient coarse-grained simulations were performed using the MARTINI 2.2 force field (46–48) with a simulation time step of 20 fs. A velocity-rescaling thermostat (modified with a stochastic term) (49) was employed with coupling constant of 1.0 ps to maintain the temperature at 310 K. Protein-lipid and solvent beads were coupled to separate baths to ensure correct temperature distribution. The pressure was kept at 1 bar using a Parrinello-Rahman barostat with a semi-isotropic coupling scheme and a coupling constant of 12 ps. All non-bonded interactions, including van der Waals forces were cut-off at 1.1 nm. The relative dielectric constant was set to 15.

Since MARTINI does not explicitly describe backbone hydrogen bonds, we imposed the secondary structures (*α*-helices) on the peptides throughout the entire simulation run. The peptide C-terminal capping was modeled by removal of the charge and changing the backbone bead type to neutral.

#### System Preparation

The membrane was assembled in the *XY* -plane using the CHARMM-GUI web server (50). The lipid bilayer was composed of 378 POPE and 126 POPG equally distributed lipids in both leaflets. Roughly 30 water beads per lipid were added (a single bead corresponds to four water molecules) and also NaCl ions were added at a concentration of 130 mM. Two P/L ratios – 1/42 and 1/21, were considered for MG2a, L18W-PGLa and MG2a/L18W-PGLa (1:1 mol/mol) placing an equal number of randomly distributed peptides into each membrane leaflet. Dimerization was derived from analyzing distances between the peptide centers of mass and the peptide termini. If at least two of these distances were smaller than 1 nm, then the peptides were considered to be in a dimer. As control we performed an unbiased simulation starting with peptide heterodimers taking all parallel/antiparallel variants of mutual peptide alignment into accoun and analyzed as a function of time. Finally, we performed a biased simulation, where MG2a and L18W-PGLa were restrained in heterodimers via a flat-bottom potential. The potential was applied to the distances between peptide centers of mass for separations larger than 1 nm with a force constant of 1000 kJ mol^−1^ nm^-2^.

## RESULTS

### Scattering Experiments

#### Membrane structure and peptide location

We determined the structural response of POPE/POPG (3:1 mol/mol) LUVs to both peptides, including their equimolar mixture (P/L = 1/200, *T* = 35°C) using a joint analysis of scattering data with a total of four contrasts (SAXS, SANS: 50, 75 100% D_2_O). No vesicle aggregation or formation of multilamellar aggregates occurred at this peptide concentration as evidenced by the pure diffuse nature of all scattering patterns (Fig. 2, Figs. S2–S4). This enabled us to derive the peptides’ effect on membrane structure in detail using the analysis described in the previous section.

**Figure 2:**
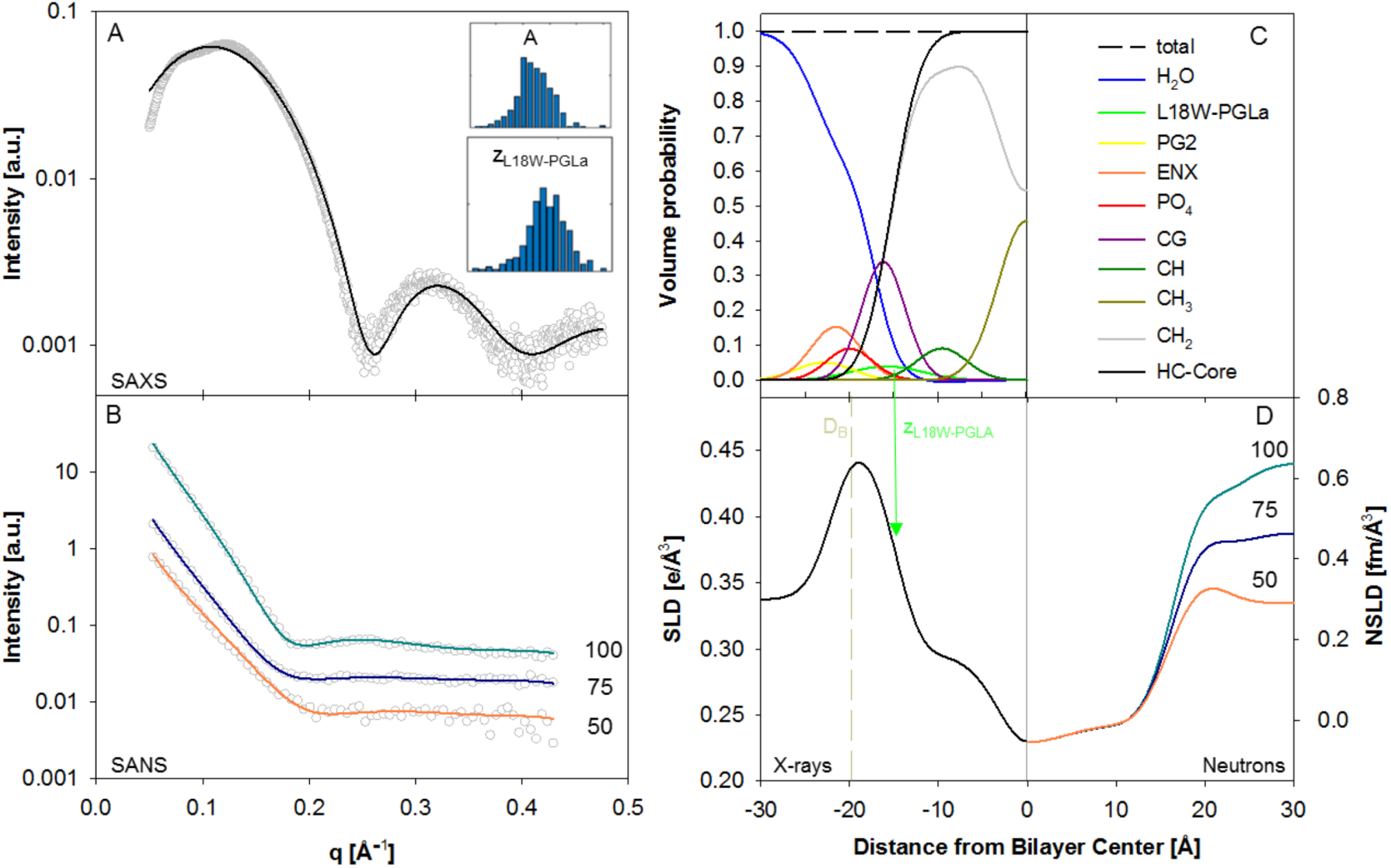
SDP analysis of POPE/POPG (3:1 mol/mol) in the presence of L18W-PGLa (P/L = 1/200). Panels (A) and (B) show the calculated model fits from averaged parameters (solid lines) for SAXS an SANS data, obtained from 400 independent optimization runs. Insets in (A) show histograms of the area per unit cell, *A*_U_, and the position of the peptide in the bilayer, *z*_P_. Panel (C) shows the volume probability distribution of the bilayer and panel (D) displays the corresponding electron and neutron scattering length densities. The arrow indicates location of *z*_P_ and the dashed-line shows the position of *D*_B_/2.

Pure POPE/POPG bilayers serve as a reference system for the present study. Our SDP analysis yields 60.56 ± 0.10 Å^2^ for the lateral area per lipid (Tab. 1), see Fig. S2, for corresponding fits and Tab. S1 for all parameter values. This value compares well to *A*_U_ = 59.8 Å^2^ obtained by molecular averaging the individual areas per lipid reported for POPE and POPG (24, 25), which supports our analysis. Moreover, our model shows a good agreement with the scattered intensities in the presence of peptides (Fig. 2), Fig. S3, S4), which further supports our assumption of evenly distributed peptides. An equal distribution of peptides in both leaflets is expected to need extended sample equilibration times, such as realized for the here presented data, during which peptides may spontaneously translocate through the membranes. We note, however, that we cannot fully exclude the presence of asymmetric transleaflet distributions of the peptides. Disentangling leaflet compositional differences is beyond the resolution of the here presented experiments.

**Table 1:** Effect of magainins on the structure of POPE/POPG (3:1 mol/mol) bilayers.

| sample | $A_U$ [$\text{\AA}^2$] | $D_B$ [$\text{\AA}$] | $D_C$ [$\text{\AA}$] | $z_P$ [$\text{\AA}$] | $D_{HD}$ [$\text{\AA}$] |
| --- | --- | --- | --- | --- | --- |
| POPE/POPG | $60.56 \pm 0.10$ | $39.15 \pm 0.06$ | $15.34 \pm 0.02$ (15.10)<sup>a</sup> | | $4.19 \pm 0.05$ |
| + L18W-PGLa | $61.26 \pm 0.13$ | $39.49 \pm 0.08$ | $15.16 \pm 0.03$ (15.06)<sup>a</sup> | $15.75 \pm 0.44$ | $4.07 \pm 0.06$ |
| + MG2a | $62.00 \pm 0.09$ | $39.17 \pm 0.06$ | $14.98 \pm 0.02$ (15.01)<sup>a</sup> | $16.29 \pm 0.50$ | $3.80 \pm 0.06$ |
| + L18W-PGLa/MG2a | $63.29 \pm 0.06$ | $38.31 \pm 0.03$ | $14.68 \pm 0.01$ (14.93)<sup>a</sup> | $15.98 \pm 0.43$ | $4.45 \pm 0.04$ |

<sup>a</sup> values in brackets are derived from all-atom MD simulations.

All peptides caused significant modulations of membrane structure (Tabs. 1) with L18W-PGLa having the least effect, e.g., the decrease of thickness of the hydrophobic core 2Δ*D*_C_ ∼ −0.4 Å and Δ*A*_U_ ∼ 0.7 Å^2^. Interestingly, *D*_B_ increased slightly. This can be mainly attributed to the increased volume of the unit cell due to contributions from L18W-PGLa (see Eq. (1)). For MG2a in turn, no significant changes were observed for *D*_B_, while 2*D*_C_ and *A*_U_ changed about twice as much compared to L18W-PGLa. In case of the equimolar peptide mixture Δ*A*_U_ ∼ 2.7 Å^2^ is most significant, leading even to a decrease of *D*_B_. Since 2Δ*D*_C_ does not contain contributions from peptide volumes, it consequently is the appropriate parameter to measure membrane thinning. For the studied peptides membrane thinning follows the order L18W-PGLa/MG2a > MG2a > L18W-PGLa. We note that the thinning observed for L18W-PGLa/MG2a cannot be explained by a simple linear combination of the thinning effects of the individual peptides. In particular the experimental form factors of POPE/POPG in the presence of L18W-PGLa and MG2a cannot be averaged to yield the form factor observed upon addition of the equimolar mixture, which should be possible if the two peptides were not interacting with each other (Fig. S5). This supports the formation of heterodimers at significantly lower P/L than previously reported (7, 9).

It is particular interesting to relate membrane thinning to the location of the peptides within the bilayer (Tab. 1). L18W-PGLa is positioned slightly below, and MG2a slightly above the glycerol backbone (CG group), respectively as observed by comparing *z*_P_ and *D*_C_. The peptide mixture behaved similar to MG2a with a preferential location just above the lipid backbone. Thus, pronounced thinning effects are observed for peptides located further away from the membrane center, as well as for L18W-PGLa/MG2a, presumably due to dimer formation.

Finally, we present results for the distance between CG and the PO_4_ groups, *D*_HD_. Because of the rotational flexibility of the lipid head group, this value is a measure of the average headgroup tilt projected on to the normal of the lipid bilayer. That is, the lowest *D*_HD_ found in the presence of MG2a indicates that the lipid headgroups are more tilted toward the membrane than for L18W-PGLa/MG2a with the largest *D*_HD_-value.

#### Effect of Temperature and Peptide Concentration

We varied the temperature in the range from 35 − 50°C to see whether the two peptides and their mixture induce specific changes to membrane structure (Figs. S6–S8). Additionally, we increased the peptide concentration ensuring that Bragg peaks resulting from vesicle aggregate changes do not dominate the scattered intensities. In the case of MG2a and L18W-PGLa this allowed us to go up to P/L = 1/50 (Figs. S7 and S8). For the 1:1 peptide mixture we observed the onset of MLV formation (indicated by a low intensity peak) already at P/L = 1/200 (Fig. S9). We thus restricted our analysis to P/L = 1/400 − 1/200 for the equimolar mixture. Peptide induced formation of coupled bilayers will be discussed in the subsequent paper.

Compared to the above discussed joint SAXS/SANS data, the performed temperature-dependent SAXS experiments provided limited structural resolution. Nevertheless relative structural changes of structural parameters such as *A*_U_ as a function of temperature can be retrieved reliably (Figs. S10, S11), using reported lipid volume temperature dependencies (24, 25). Figure 3 shows the thermal expansion of the area per unit cell for POPE/POPG bilayers (Δ*A*_U_/Δ*T*), for the presently studied magainins as function of peptide concentration resulting from this analysis. In general we find that the area expansion decreases with peptide concentration and levels off at about 0.22Å^2^/K, independent of the specific structure of the added peptides including their mixture. This shows that membranes are increasingly less able to expand laterally with increasing temperature as a result of the peptide induced membrane perturbation.

**Figure 3:**
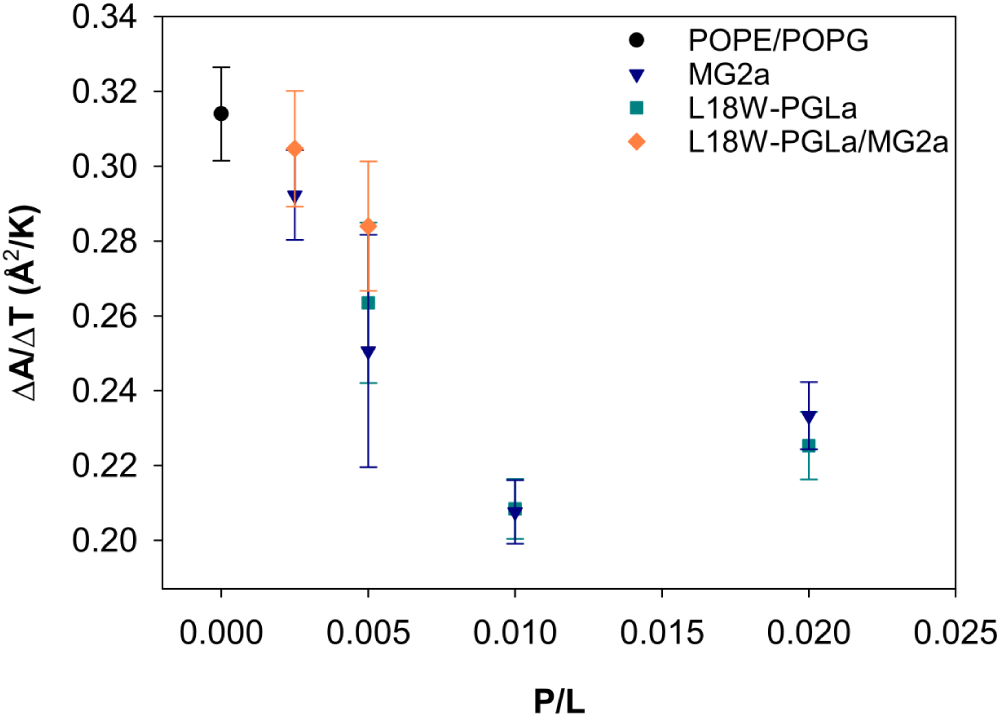
Dependence of the thermal area expansion coefficient of POPE/POPG (3:1 mol/mol) bilayers on peptide concentration (circles: pure bilayer, triangles: MG2a, squares: L18W-PGLa, diamonds: L18W-PGLa/MG2a (1:1 mol/mol)).

### All-atom and coarse-grained simulations

To gain additional insights on the molecular level, we performed MD simulations of both peptides within POPE/POPG (3:1 mol/mol) bilayers. Figure S12 compares the form factors derived from all-atom MD simulations to experimental data. The remarkable agreement in particular for *q* > 0.15 Å provides a valuable validation of the results presented below.

#### Individual peptides

Firstly, we performed all-atom simulations with peptide monomers, where each membrane leaflet contained a single peptide to ensure membrane and system symmetry. Throughout the simulations, both peptides retained their mostly *α*-helical conformation and remained oriented parallel with respect to the membrane plane (Fig. 4). The exception was MG2a, which showed a partial loss of helicity at the C-terminus in one leaflet. Figure 4 shows snapshots from the end of the 500 ns long simulations together with the depth of peptide insertion. L18W-PGLa was found to be slightly deeper in the headgroup region compared to MG2, which is consistent with our SANS/SAXS analysis.

**Figure 4:**
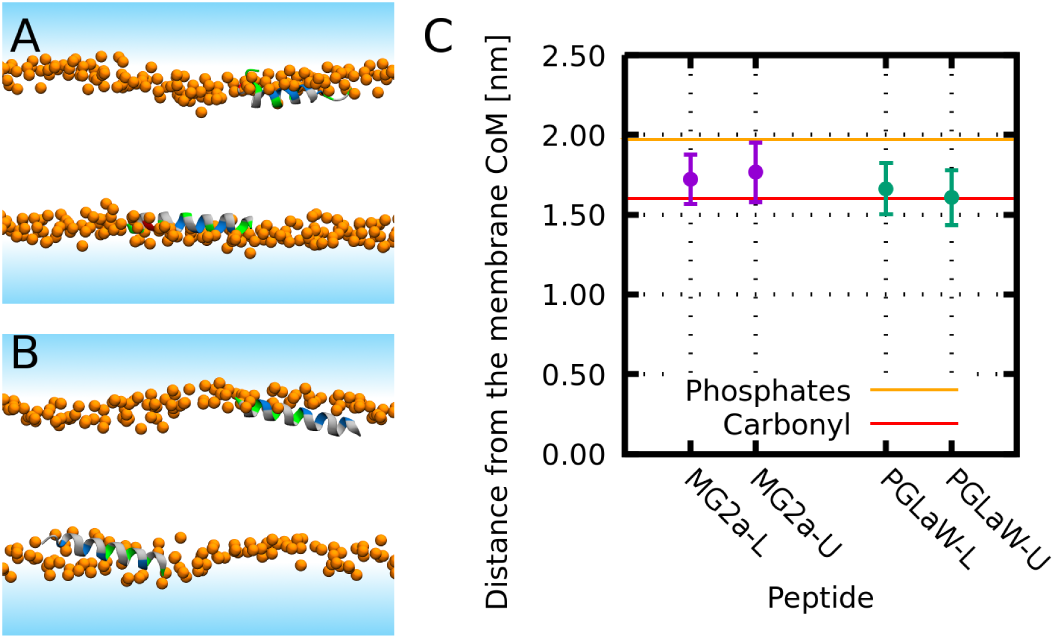
Membrane positioning of MG2a and L18W-PGLa peptides. Left side shows the snapshots from the end of 500 ns long MD simulations of (A) MG2a and (B) L18W-PGLa adsorbed at the membrane surface. Panel C shows the peptide positions in both leaflets averaged over trajectories. Snapshots color coding: Lipid phosphate atoms are shown as orange spheres. Solvent is represented by a blue-shaded area and lipid tails are not shown for clarity. The peptide secondary structure is shown in a cartoon representation, colored by residue type (nonpolar: gray, polar: green, acidic: red, and basic: blue).

We observed a local membrane modification in the vicinity of the peptides in agreement with our scattering data analysis. Lipids changed their tilt and conformation in order to fill the hydrophobic void below the inserted peptide, as shown previously (51). To quantify this effect, we calculated the density distribution of the methyl groups of the lipid tails in the peptide aligned trajectories (Fig. 5 *A, B*). The methyl density was locally increased below the peptides filling the available space between peptide side-chains. The consequent effects on hydrocarbon chain packing are observed in the order parameter profiles dependence as a function of the distance from the peptides Fig.6 *A, B* (see also Figs. S13, S14). In general, our analysis showed that palmitoyl chains are mostly affected at intermediate segments, whereas oleoyl hydrocarbons experience most significant changes toward the hydrocarbon tails. Moreover, MG2a appears to induce a slight increase of order close to the POPG glycerol backbone of the unsaturated and a more pronounced disordering of its saturated hydrocarbon chain, respectively. Overall, the effects on hydrocarbon packing lead to membrane thinning in good agreement with our scattering data analysis (see Tab. 1). The changes in density distributions for individual groups of membrane are depicted in Figs. S15 and S16.

**Figure 5:**
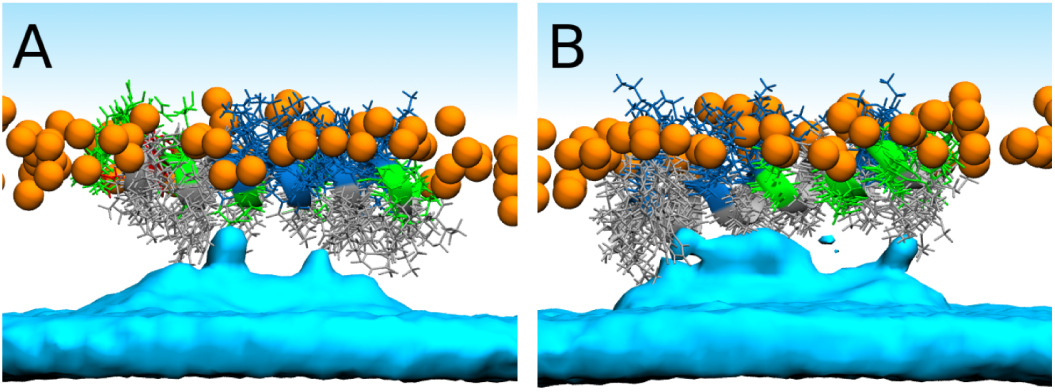
Averaged densities of terminal hydrocarbon CH_3_ groups (cyan surfaces) after insertion of MG2a (A) and L18W-PGLa (B) in POPE/POPG bilayers.

**Figure 6:**
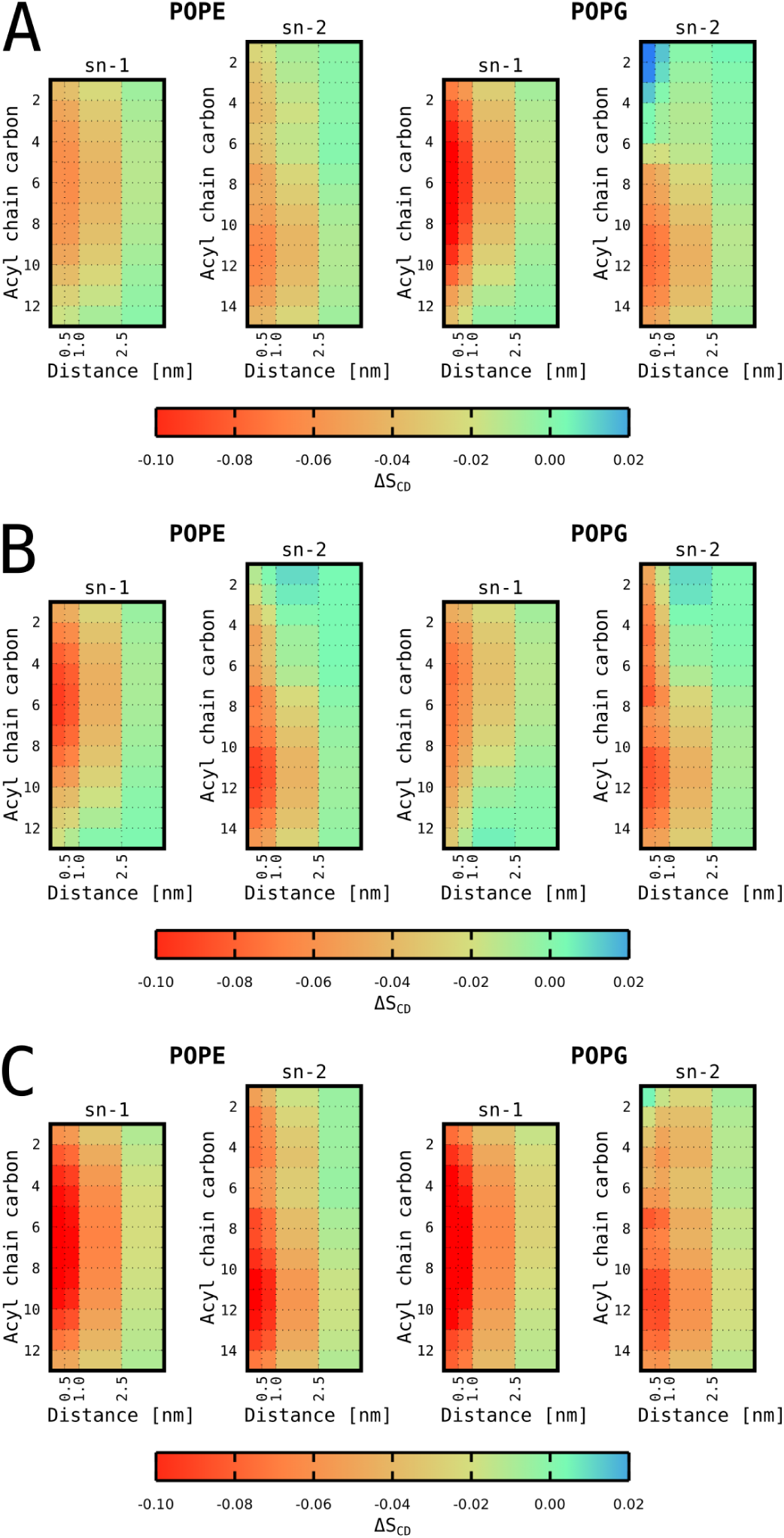
Changes of the lipid tail order parameters as a function of distance from the peptide (A) MG2a, (B) L18W-PGLa, and (C) MG2a+L18W-PGLa.

### Effects of peptide dimers

MG2a and PGLa peptides were previously shown to prefer parallel dimers using a coarse-grained model (52), which also agrees with experimental findings (7, 9). We verified this tendency also for our bacterial membrane mimic. Due to the low number of dimer formation events at low peptide concentrations within the length of our simulations (20 µs), we had to increase P/L ratios from 1/42 to 1/21 in order to facilitate the analysis. We note, however, that this did not lead to the formation of a transmembrane pore as observed in simulations using dilaureoylphosphatidylcholine bilayers (52). This is due to the larger membrane thickness and tighter interfacial lipid packing of POPE/POPG (3:1 mol/mol) bilayers (5) and allowed us to derive the dimerization behavior even at elevated peptide levels. MG2a/L18W-PGLa mixtures starting from random configuration showed the strongest preference for dimerization, followed by MG2a and L18W-PGLa (Fig. 7). MG2a/L18W-PGLa mixtures mainly formed parallel heterodimers, while MG2a and L18W-PGLa preferentially formed anti-parallel homodimers. An additional independent 40 µs long coarse-grained simulations starting with peptides preformed in various heterodimer configurations further corroborated the higher stability of parallel heterodimers as compared to other peptide-peptide alignments (Fig. S20).

**Figure 7:**
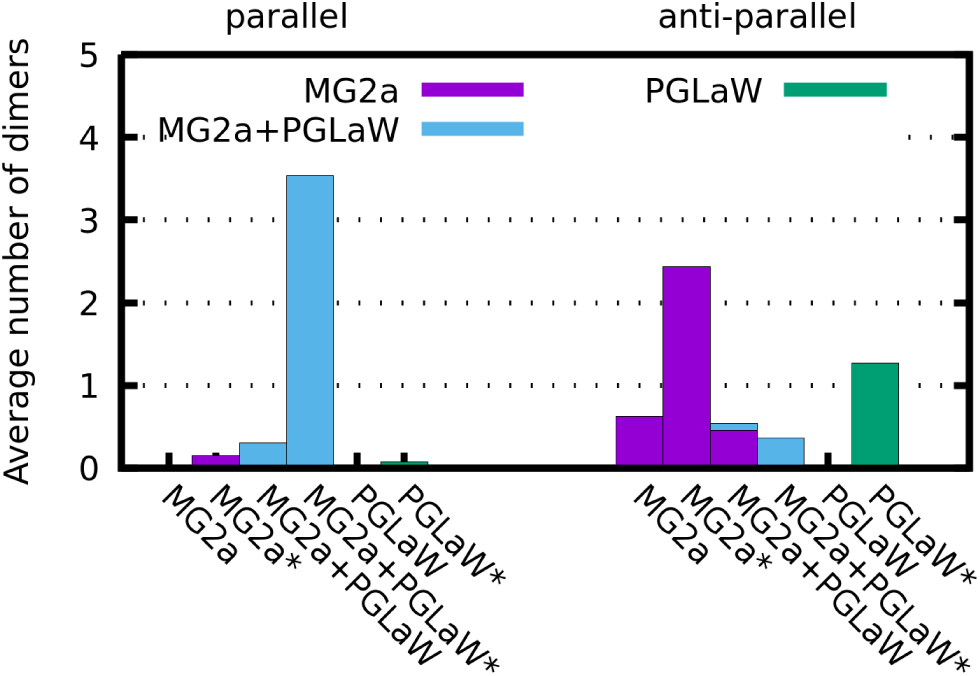
Time averaged number of dimers formed in all simulated coarse-grained systems. Left group corresponds to parallel dimers, while anti-parallel dimers are in the right group. Systems were simulated at either 1/42 or 1/21 (denoted by asterisk) peptide-to-lipid ratio. Purple bars represent MG2a homodimers, L18W-PGLa homodimers are green, and MG2a+L18-WPGLa heterodimers are shown in cyan.

Since the peptide-peptide interactions might be overestimated in coarse-grained simulations (53), we performed additional all-atom simulations starting from preformed dimers. In agreement with our coarse-grained simulations, we found that parallel heterodimers were stable on the simulated timescale.

Based on the preference of MG2a/L18W-PGLa mixtures to form parallel MG2a/L18W-PGLa heterodimers, we investigated the effect of this heterodimer on membrane structure using all-atom simulations of parallel heterodimers in both membrane leaflets (see Fig. S17 for final snapshot of the 500 ns trajectory). In agreement with experimental data heterodimers lead to increased membrane thinning (Tab. 1) and significantly pronounced hydrocarbon chain packing defects (Fig. 6 *C*) as compared to the individual peptides. This supports our above notion that the experimentally observed enhanced membrane perturbation is a result of L18W-PGLa/MG2a dimer formation at low peptide concentrations.

In the next step we interrogated our all-atom simulations for the effect of dimerization on peptide location within bilayers. Compared to MG2a and L18W-PGLa (Fig. 4 *C*), L18W-PGLa/MG2a heterodimers inserted more shallow into the membrane (Fig. S18). An analysis of our coarse-grained Martini simulations at both peptide concentrations showed that the transbilayer position of the heterodimers was within uncertainty almost equal to that of MG2a (Fig. 8, Fig. S19). In order to see, whether the dynamics of dimer formation might affect their insertion depth, we artificially kept the peptides in parallel heterodimer configuration in an additional simulation (for details see the Methods section). This simulation decreased the uncertainty of the peptide position significantly, but gave the overall same result for the dimer position as the unconstrained coarse grained simulations, i.e. dimers are located closely to the phosphate group. The main difference compared to all-atom simulations is the slightly less deeper insertion of MG2a, which could be due to force field issues. However, on an absolute scale these differences are only minor. Moreover, results of both simulation models even agree considering the positional uncertainty of the peptides (Fig. 8, Figs. S18, S19). Regarding experimental data (Tab. 1), we find an overall reasonable agreement, with respect to the peptide position. Specifically, there is broad consensus between the here applied different experimental and computational techniques that L18W-PGLa positions itself further out from the bilayer center, when dimerizing with MG2a.

**Figure 8:**
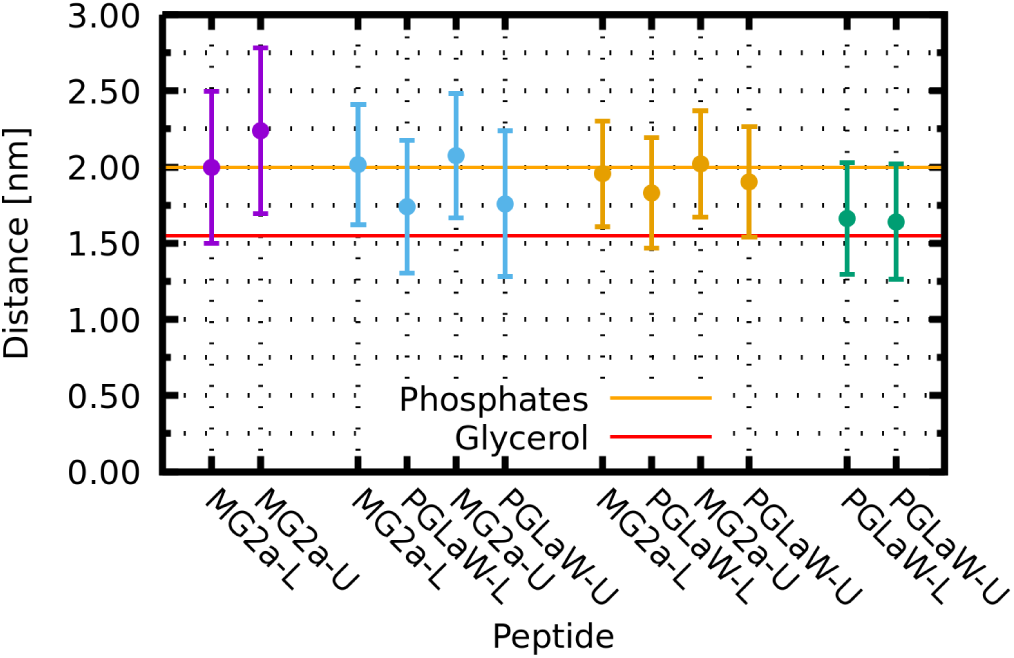
Effect of peptide dimerization on the insertion depth of peptides in POPE/POPG bilayers from coarse-grained simulations. Data show the distances of the peptides from the membrane’s center of mass for the individual peptides (MG2a (purple), L18W-PGLa (green)), as well as unconstrained (blue) and constrained (orange) L18W-PGLa/MG2a (P/L = 1/21). The distances were averaged over 20 µs for all peptides and are presented for (L)ower and (U)pper membrane leaflets individually. Error bars represent the standard deviation. Orange and red horizontal lines represent the approximate position of phosphate and glycerol groups.

## DISCUSSION

We combined SAXS/SANS experiments with all-atom and coarse-grained MD simulations to interrogate mutual interactions between L18W-PGLa, MG2a, and fully hydrated POPE/POPG (3:1 mol/mol) bilayers. The focus of the present work is on low peptide concentrations, i.e. where equimolar mixtures of both peptide do not permeabilize POPE/POPG bilayers synergistically (5). Yet, we demonstrate that the peptide mixture leads to significantly enhanced membrane perturbations even at low concentrations.

We developed a SDP analysis for peptide containing lipid membranes, which allowed us to determine the peptide position in the bilayer with high accuracy at low peptide concentrations by a simultaneous statistical analysis of four differently contrasted SAXS/SANS experiments. The observables were in excellent agreement with our MD simulations, enabling additional insight on lipid-peptide and peptide-peptide interactions. Note that previous reports on similar systems using solid state ^15^N-NMR were not sensitive to the penetration depth of peptides within the headgroups. Both our experimental and simulation data consistently showed that MG2a inserts slightly less into the bilayer than L18W-PGLa (Tab. 1, Fig. 4 *C*). This can be understood in terms of the larger hydrophilic surface of MG2a compared to L18W-PGLa due the larger number of polar residues (5).

In agreement with ^15^N-NMR data on POPE/POPG (3:1 mol/mol) (13) and POPE-enriched bilayers (11) we found that both peptides adopt a surface-aligned topology, even in the case of equimolar L18W-PGLa/MG2a mixtures. The surface-aligned topology caused significant disorder of hydrocarbon chain packing, which was most pronounced in the vicinity of the peptides. In particular, we observed that the methyl termini of the hydrocarbons filled large fractions of the void below the peptide (Fig. 5), consistent with previous reports on dimple formation and membrane thinning (54). Membrane thinning was observed for both peptides (Tab. 1), consistent with a surface-aligned topology. The pronounced membrane thinning for equimolar mixtures of the two peptides provides indirect evidence for dimer formation. This is further supported by our experimental form factors, which were determined from a statistical data analysis and which clearly show that the effects of the individual peptides cannot be simply combined to yield the membrane structure in the presence of both peptides (Fig. S5). Hence, PGLa and MG2a seem to form dimers at concentrations much lower than reported previously (7, 9, 52).

Using all-atom MD simulations we derived order parameter profiles as a function of distance from the peptides. In agreement with a previous NMR study (13) both peptides were found to perturb the saturated hydrocarbons of surrounding lipids (Fig. 6). Our results suggest that MG2a is more effective than L18W-PGLa in doing so, which correlates with the further outward location of MG2a within the membrane and its increased bulkiness (molecular volumes: *V*_L18W-PGLa_ = 4927.8 Å^3^, and *V*_MG2a_ = 5748.0 Å^3^). MD simulations additionally allowed us to probe order parameter profiles of unsaturated hydrocarbons. Interestingly, we found that MG2a increased the order of the oleoyl chain close to the glycerol backbone of POPG only, while a general decrease of this chain’s order parameter was observed for POPE at all segments for both peptides (Fig. 6 *A, B*, Figs. S13, S14). This indicates specific interactions of MG2a with POPG, which are located around the positively charged amino acids as revealed a detailed analysis of MD data (Fig. S21). Finally, assuming heterodimers, all effects observed with respect to the order of hydrocarbon chains were significantly more pronounced as compared to single peptides (Fig. 6 *C*), in agreement with our experimentally observed changes of membrane structure. This is also consistent with previous reports (see, e.g. (55)), showing that the formation of dimers amplifies the perturbation of the lipid tails in the vicinity of peptide dimer as compared to monomers.

Our simulations indicated that homodimers formed preferentially in antiparallel configuration, while L18W-PGLa/MG2a formed mainly parallel heterodimers (Fig. 7), which is in agreement with a previous study (52). Interestingly, heterodimer formation has been reported for significantly different lipid bilayers. Thus, dimerization of PGLa and MG2a, does not appear to be highly specific to lipid composition, although we cannot comment on the onset of dimerization in other lipid bilayers from the present study. The analysis of enthalpic interactions between different amino acids showed that dimer formation is stabilized by salt bridges between MG2a-Glu19 and Lys12 or Lys15 residues of L18W-PGLa (Fig. S22) in agreement with (52). Moreover, there are significant hydrophobic interactions between the peptides. For example, Ulmschneider *et al.* reported a stabilization of PGLa homodimers by Gly/Ala interactions at high P/L ratios (56). It appears, however, that the sum of all these interactions leads to a preferential formation of L18W-PGLa/MG2a heterodimers as compared to L18W-PGLa/L18W-PGLa or MG2a/MG2a homodimers, which is consistent with previous observations (6).

Most interestingly, peptide dimerization affects the penetration depth of the surface-aligned peptides. In particular L18W-PGLa moves somewhat further out in the headgroup region, when it associates with MG2a (Tab. 1, Fig. 8, Figs. S18, S19). This leads to a larger void created in the membrane interior just below the peptides, causing an increased change of membrane structure. Hence, the ability to form a dimers appears to be important for the membrane perturbation efficacy of the studied peptides.

## CONCLUSION

Our study suggests that an early onset of the formation of peptide dimers is the key event to the enhanced activity of L18W-PGLa/MG2a mixtures. Previously, we speculated that L18W-PGLa causes a deeper insertion of MG2a into the bilayer (5). Indeed, the present work shows the opposite, i.e. that L18W-PGLa moves further out from the membrane center when forming a heterodimer with MG2a. These heterodimers perturb membranes significantly more than the sum of the effects induced by individual (non-interacting) peptides. Apparently, this ‘synergistic’ dimerization is not sufficient to allow enhanced leakage of dyes (although smaller polar molecules might already permeate the bilayer) (5). For membrane leakage to occur, higher peptide concentrations are required. The corresponding membrane restructuring effects will be described in the subsequent paper.

## SUPPORTING MATERIAL

Supporting Materials and Methods, twenty three figures, and one table are available at …

## AUTHOR CONTRIBUTIONS

M.P. performed the experimental research, analyzed the data, and wrote the article. I.K. carried out all simulations, analyzed the data, and wrote the article. M.-S.A. performed SANS experiments. R.V., K.L. and G.P. designed the research and wrote the article.

## ACKNOWLEDGMENTS

This work was supported by the Austrian Science Funds FWF (project No. I1763-B21 to K.L.), the Czech Science Foundation (grant 17-11571S to R.V.) and the CEITEC 2020 (LQ1601) project with financial contribution made by the Ministry of Education, Youths and Sports of the Czech Republic within special support paid from the National Programme for Sustainability II funds. Computational resources were provided by the CESNET LM2015042 and the CERIT Scientific Cloud LM2015085, provided under the programme ‘Projects of Large Research, Development, and Innovations Infrastructures’. This work was supported by The Ministry of Education, Youth and Sports from the Large Infrastructures for Research, Experimental Development and Innovations project ‘IT4Innovations National Supercomputing Center – LM2015070’. We acknowledge SOLEIL for provision of synchrotron radiation facilities and we would like to thank Javier Perez for assistance in using beamline SWING. This work is based upon experiments performed at the KWS-1 instrument operated by JCNS at the Heinz Maier-Leibnitz Zentrum (MLZ), Garching, Germany.

## SUPPORTING CITATIONS

Reference (57) appears in the Supporting Material.

## Notes

#### Summary of Updates

In particular, we have rewritten completely the simulation sections on peptide dimerization and also performed an additional coarse grained simulation to check for the stability of the different dimer concentrations (Fig. S20). The results confirmed our previous conclusion that parallel PGLa-MG2a heterodimers are most stable. Further, we also carefully clarify our statements concerning synergy, motivating the study at low peptide concentrations. Synergistic activity of peptides on lipid model systems is frequently attributed to the increased leakage of fluorescent dyes of a specific size and has also been done here. However, this does not preclude any synergistic interactions between the peptides at much lower concentrations, which just is not perturbating enough the membrane to allow for the passage of the dyes. In the present case we demonstrate the association of the peptides in form of parallel heterodimers already at low concentration. The evidence is mainly based on SAXS/SANS data and possible only through the combination of 4 different contrasts with a statistical global fitting procedure. This is also distinct and superior to previous similar approaches. Note that the experimental form factors of the system in the presence of the single peptides cannot be combined linearly (as should be possible if the peptides are not interacting) to give the form factor in the presence of the peptide mixture (new Fig. S5).

