## Supporting Material for "Magainin 2 and PGLa in Bacterial Membrane Mimics I: Peptide-Peptide and Lipid-Peptide Interactions"

*M.P., I.K., M.-S.A., K.L., R.V., G. P.*

### Scattering Density Profile Model for a symmetric Membrane

POPE and POPG were parsed into $CH_3$, $CH_2$, CH, CG, $PO_4$, ENX and PG2 groups as reported previously (1, 2) (Fig. S1). Since we studied POPE/POPG mixtures the several of these groups were combined in order to reduce the number of adjustable parameters. In particular, we merged the $CH_3$, $CH_2$, CH, CG, $PO_4$ groups and fitted only ENX and PG2 individually. Except for $CH_2$ all lipid groups were modelled with Gaussians

$$P_i^X(z) = S_i^X \frac{n_i V_i}{A_{\mathrm{U}} \sigma_i} \left[ e^{-\frac{(z+z_i)^2}{2\sigma^2}} + e^{-\frac{(z-z_i)^2}{2\sigma^2}} \right] \tag{S1}$$

of width $\sigma_i$ with $i \in \{\mathrm{CH_3, CH, CG, PO_4, ENX\ and\ PG2}\}$, where $V_i$ is the volume of each group and $n_i$ is number of type $i$ components (e.g. $n_{\mathrm{CH_3}} = 2$).

NMR experiments demonstrated that L18W-PGLa and MG2a align parallel to the membrane surface in POPE/POPG bilayers (3). Their volume distribution functions were therefore described analogously to Eq. (S1). The scaling factor $S_i^X$ ($X \in \{\mathrm{L, P}\}$) accounts for the fraction of lipid

$$S_i^L = \frac{n_i^L}{\sum_i^n n_i^L} \tag{S2}$$

and peptide per unit cell

$$S_i^P = \frac{n_i^P}{\sum_i^n n_i^L}, \tag{S3}$$

respectively.

The overall hydrocarbon core of the unit cell was described with error functions

$$P_{\mathrm{HC}}^L(z) = \frac{1}{2} \left[ \frac{2}{\sqrt{\pi}} \int_0^{\frac{z+z_i}{\sqrt{2}\sigma}} e^{-x^2} dx + \frac{2}{\sqrt{\pi}} \int_0^{\frac{z-z_i}{\sqrt{2}\sigma}} e^{-x^2} dx \right], \tag{S4}$$

from which the $CH_2$ distribution is obtained via

$$P_{CH_2}^L(z) = P_{HC}^L(z) - P_{CH3}^L(z) - P_{CH}^L(z). \tag{S5}$$

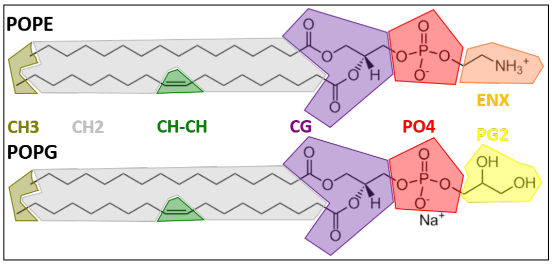


Figure S1: Schematic of POPE and POPG parsing into quasimolecular fragments. The chains consist of terminal methyl ($CH_3$), methylene ($CH_2$) and methine (CH), whereas the headgroup region is composed of carbonyl+glycerol (CG), phosphate ($PO_4$), ethanolamine (ENX) and glycerol (PG2).

Assuming ideal volume filling at every position $z$ along the bilayer normal, we obtain the water distributions by using

$$P_W(z) = 1 - \sum_k P(z)_k, \tag{S6}$$

with $k \in \{CH_3, CH_2, CH, CG, PO_4, ENX, PG2, \text{and } P\}$.

Finally, the neutron and X-ray form factors were calculated through

$$F(q) = 2 \int_0^{\frac{D}{2}} \Delta\rho(z) cos(qz) dz, \tag{S7}$$

where

$$\Delta\rho(z) = \sum_k \left[\rho_k(z) - \rho_W\right] P_k(z), \tag{S8}$$

is the scattering length density (SLD) contrast with respect to water $\rho_W$ and D/2 refers to any position outside the bilayer for which $\Delta\rho$(z) = 0.

The total volumes of the individual components needs to be supplied to the analysis. For example, the volume of the unit cell containing lipid and peptide

$$V_{LP} = \frac{n_{POPE} V_{POPE}}{n_{POPE} + n_{POPG}} + \frac{n_{POPG} V_{POPG}}{n_{POPE} + n_{POPG}} + \frac{n_P V_P}{n_{POPE} + n_{POPG}}, \tag{S9}$$

where $n_{POPE}$, $n_{POPG}$ and $n_P$ are the total numbers of POPE, POPG and peptide molecules, and $V_{POPE}$ = 1175.1 Å$^3$, $V_{POPG}$ = 1216.55 Å$^3$ (1, 2) are the corresponding volumes. The peptide volumes $V_P$ = 4927.8 Å$^3$ for L18W-PGLa and 5748.0 Å$^3$ for MG2a were obtained from MD simulations.

The scattered intensity is then given by

$$I(q) = \frac{K}{q^2} F(q)^2 + I_{inc}, \tag{S10}$$

where $I_{inc}$ is the incoherent background and $K$ is the instrumental scaling constant.

SAXS/SANS data were jointly fitted with a combined cost function

$$\chi^2 = \sum_{i,j} \left( \frac{I_i^j - I_{\mathrm{fit_i}}^j}{w_i^j \sigma_i^j} \right)^2, \tag{S11}$$

where $I_i^j$ is the recorded intensity of contrast j (e.g. SAXS or SANS including differently contrasted samples), $I_{\mathrm{fit_i}}^j$ is the calculated intensity, and $\sigma_i^j$ corresponds to the experimental error of the measured intensities. Additionally, specific weighting schemes were employed to account for the importance of the first minimum as well as the intensity modulations at high $q$. This was achieved by introducing weighting factors $w_i^j$ in order to decrease experimental uncertainties by in specific $q$-regions of the scattering data.

To ensure connectivity of the lipid structure and to further reduce the number of adjustable parameters, we coupled the CG group to the boundary of the hydrocarbon core analogously to a previous report (4), $z_{\mathrm{CG}} = D_{\mathrm{C}} + 1$ Å, and fixed relative distances between $z_{\mathrm{ENX}} - z_{\mathrm{PO_4}} = 1.57$ Å and $z_{\mathrm{PG2}} - z_{\mathrm{PO_4}} = 2.92$ Å, as reported in (1, 2). By definition, the position of the $CH_2$ group is constrained by $z_{\mathrm{CH_2}} = V_{\mathrm{HC}}/A_{\mathrm{U}}$, thus leaving only $z_{\mathrm{CH_3}}$, $z_{\mathrm{CH}}$, $z_{\mathrm{PO_4}}$ and the position of the peptide $z_{\mathrm{P}}$, as well as the corresponding widths as freely adjustable parameters. In cases, where only SAXS data was available a quadratic penalty function for negative water probabilities was utilized, which were caused by inappropriate positioning of the peptides. The corresponding prefactor was chosen to be as small as possible, in order to prevent a trapping of the fitting routine in a local minimum.

### Supplementary Figures

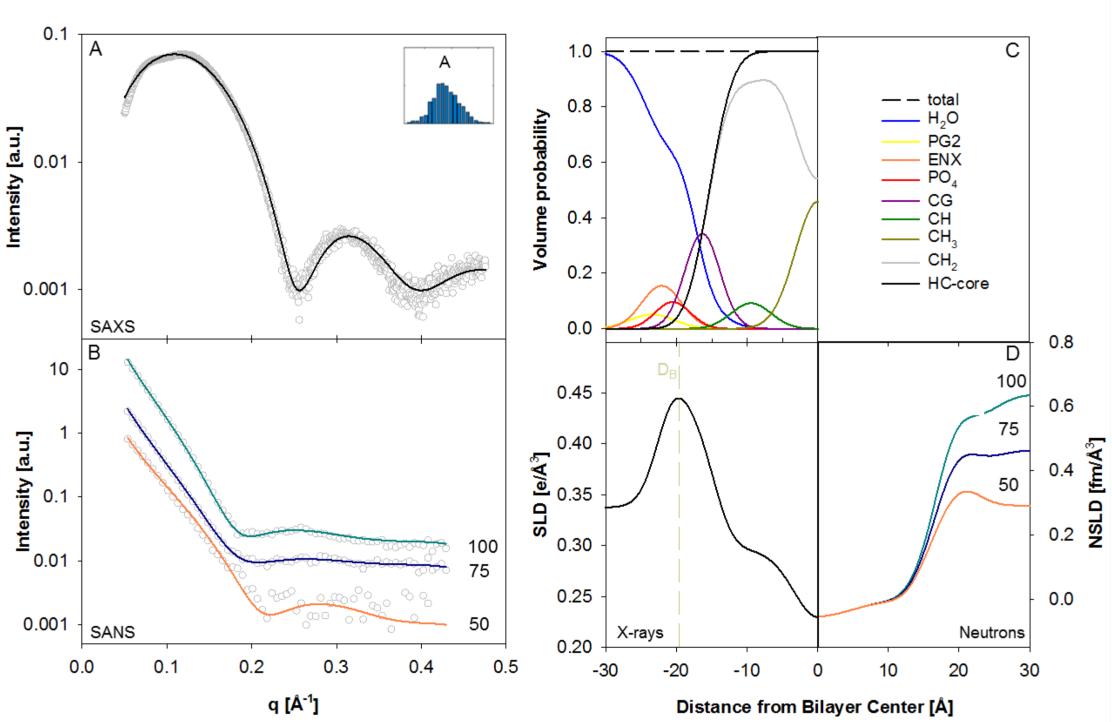


Figure S2: Joint analysis of SAXS and SANS data of LUVs (size ∼ 100 nm) composed POPE/POPG (3:1 mol/mol) at 35°C. Panels (A) and (B) show the fits (solid lines) of the joint analysis. The insert to (A) shows a histogram of the area per unit cell obtained from the statistical analysis. Panel (C) shows the volume probability distribution of the bilayer and panel (D) displays the corresponding electron and neutron scattering length densities. The Luzzati thickness, $D_{\mathrm{B}}$, is marked in the electron density profile.

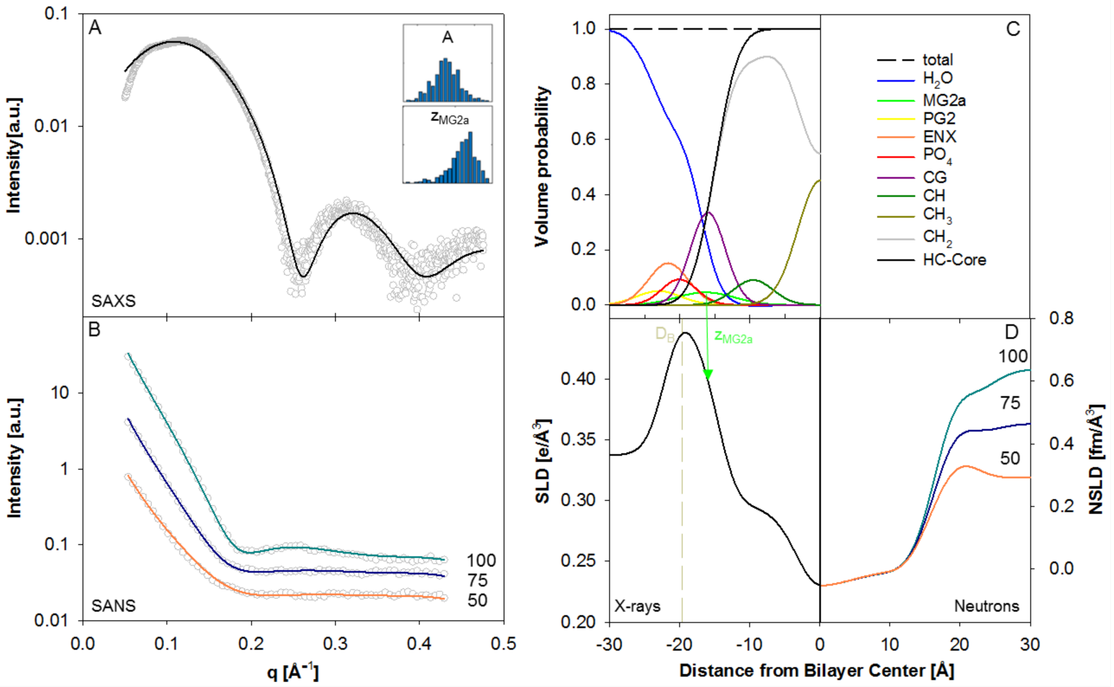


Figure S3: Joint analysis of SAXS and SANS data of liposomes composed POPE/POPG (3:1 mol/mol) in the presence of MG2a (P/L = 1/200) at 35°C. Panels (A) and (B) show the fits (solid lines) of the joint analysis. Inserts to (A) show histograms of the area per unit cell and the position of the peptide in the bilayer as obtained from the statistical data analysis. Panel (C) shows the volume probability distribution of the bilayer and panel (D) displays the corresponding electron and neutron scattering length densities. The Luzzati thickness, $D_B$, and the transbilayer position of MG2a, $z_{MG2a}$, are marked in the electron density profile.

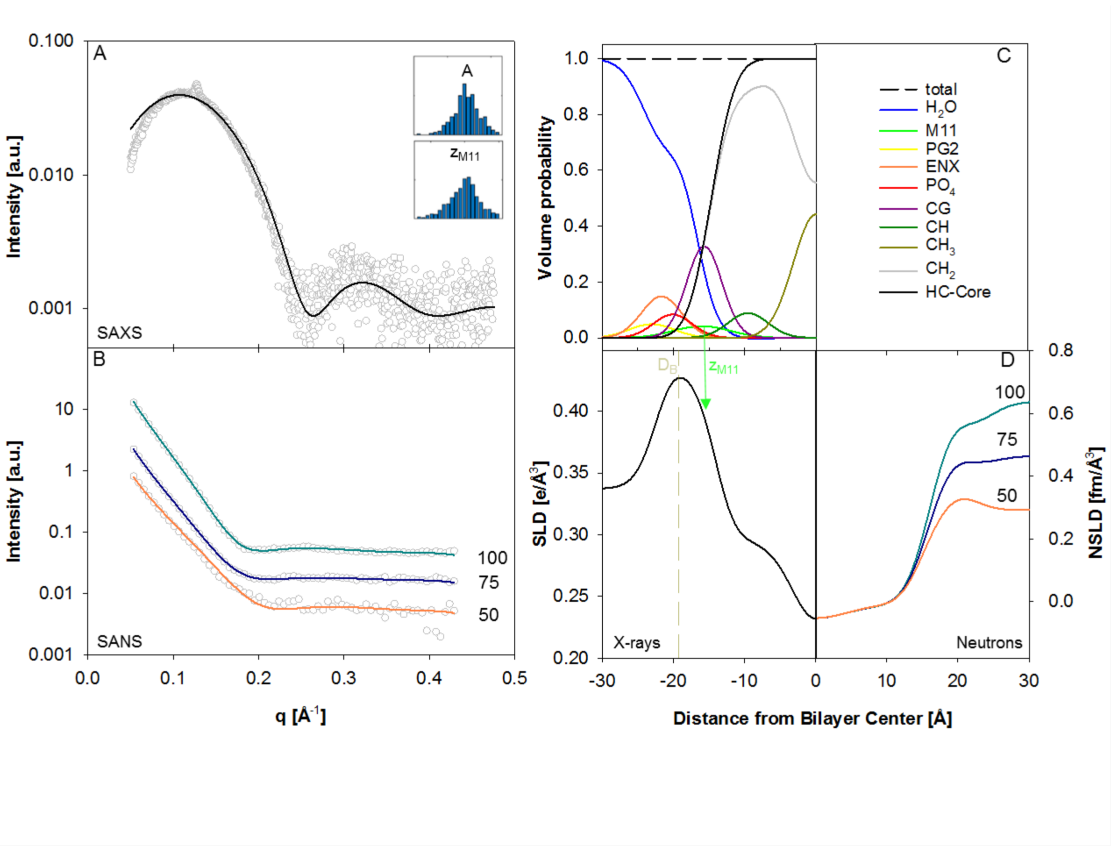


Figure S4: Joint analysis of SAXS and SANS data of liposomes composed POPE/POPG (3:1 mol/mol) in the presence of a equimolar mixture of L18W-PGLa and MG2a (M11) (P/L = 1/200) at 35°C. Panel (A) and (B) show the fits (solid lines) of the joint analysis. Inserts to (A) show histograms of the area per unit cell and the position of the peptide in the bilayer as obtained from the statistical data analysis. Panel (C) shows the volume probability distribution of the bilayer and panel (D) displays the corresponding electron and neutron scattering length densities. The Luzzati thickness, $D_{\mathrm{B}}$, and the average transbilayer peptide position, $z_{\mathrm{M11}}$, are marked in the electron density profile.

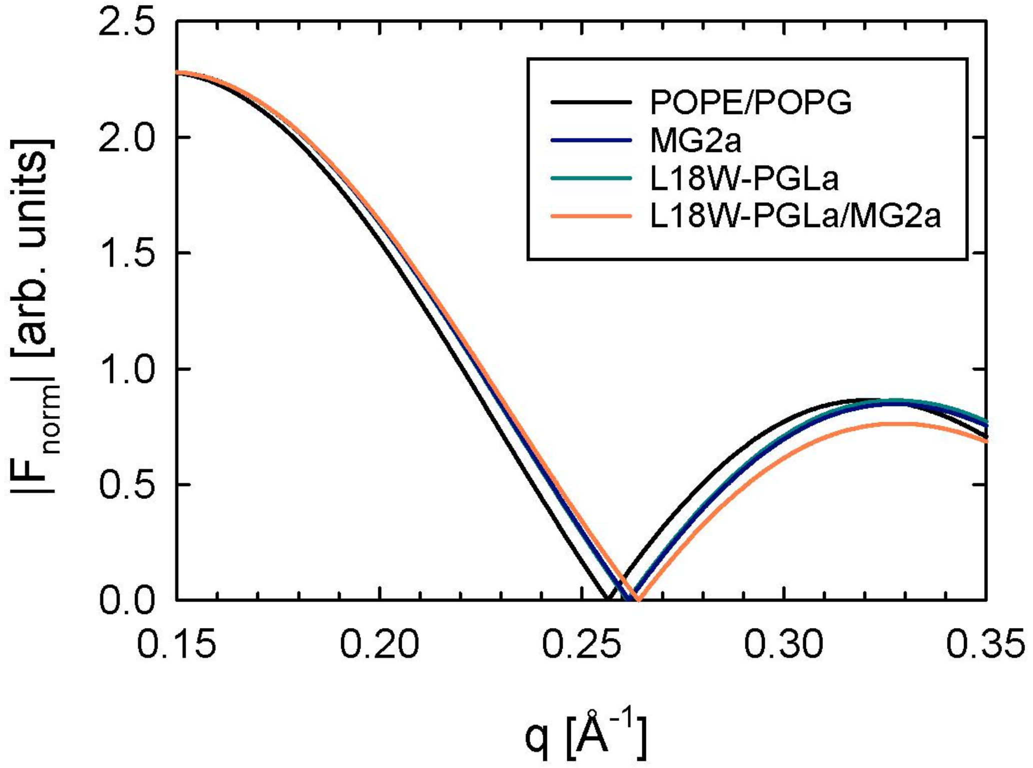


Figure S5: Normalized experimental SAXS form factors for POPE/POPG in the absence and presence of the different peptides obtained from statistical data fitting.

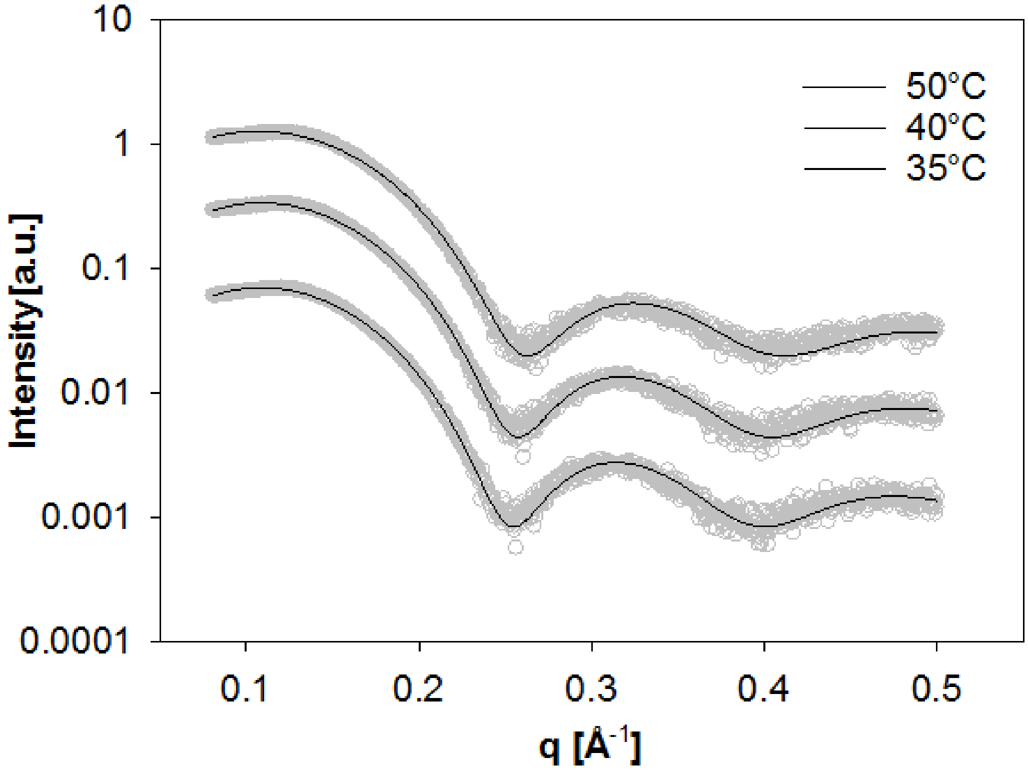


Figure S6: SAXS data of 100 nm unilamellar vesicles composed of POPE/POPG (3:1 mol/mol, open circles) at 35, 40 and 50°C. Solid lines correspond to fits with the SDP model.

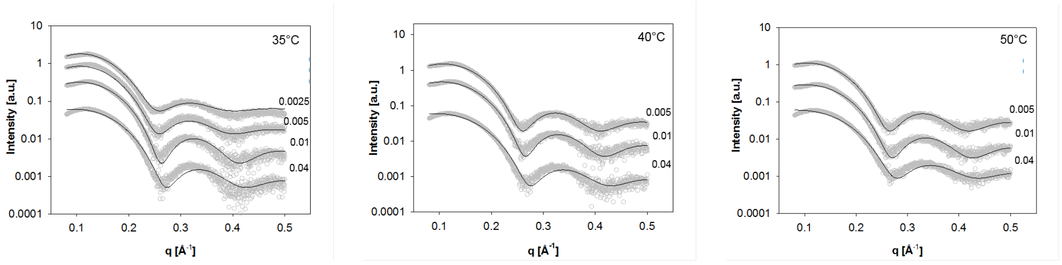


(a) (b) (c)

Figure S7: SAXS data of 100 nm unilamellar vesicles composed of POPE/POPG (3:1 mol/mol) in the presence of L18W-PGLa as a function temperature and P/L (numbers right to the scattering data). Solid lines correspond to fits.

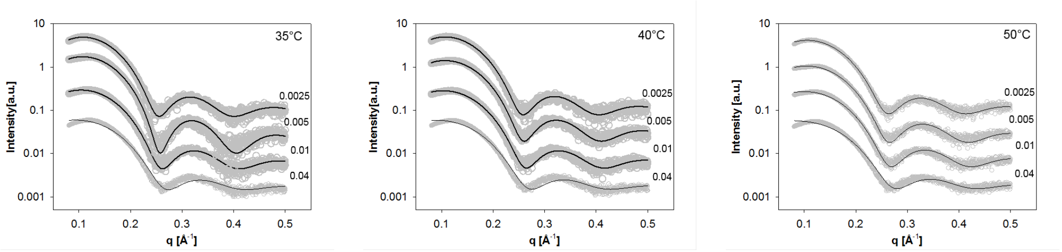


(a) (b) (c)

Figure S8: SAXS data of 100 nm unilamellar vesicles composed of POPE/POPG (3:1 mol/mol) in the presence of MG2a as a function temperature and P/L (numbers right to the scattering data). Solid lines correspond to fits.

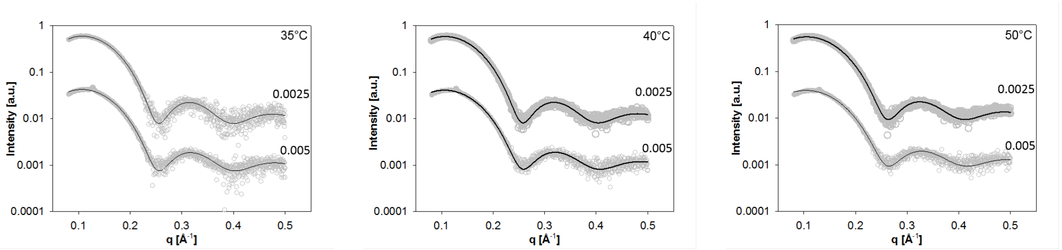


(a) (b) (c)

Figure S9: SAXS data of 100 nm unilamellar vesicles composed of POPE/POPG (3:1 mol/mol) in the presence of a equimolar mixture of L18W-PGLa and MG2a as a function temperature and P/L (numbers right to the scattering data). Solid lines correspond to fits.

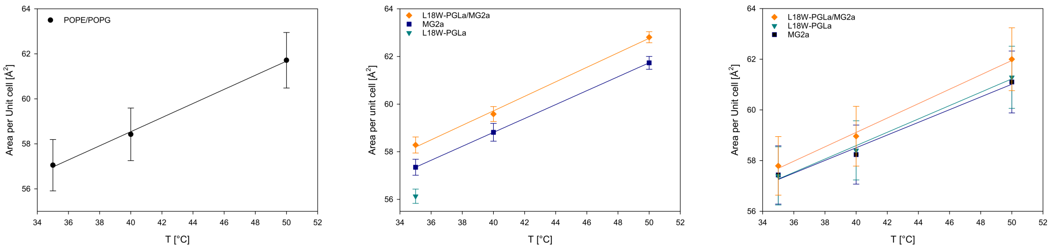


(a) (b) (c)

Figure S10: Temperature dependencies of the area per unit cell, in the absence of peptide (a) and for P/L = 1/400 (b) and 1/200 (c). Solid lines represent the fits.

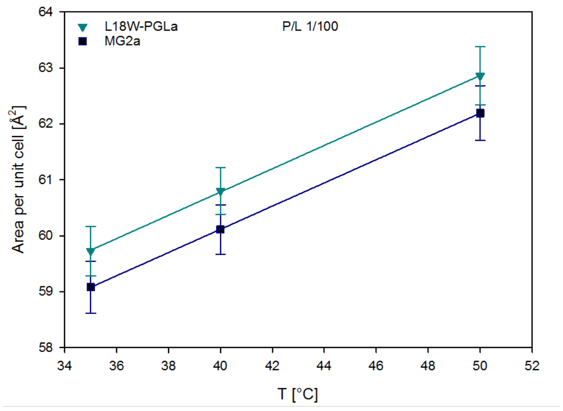


(a)

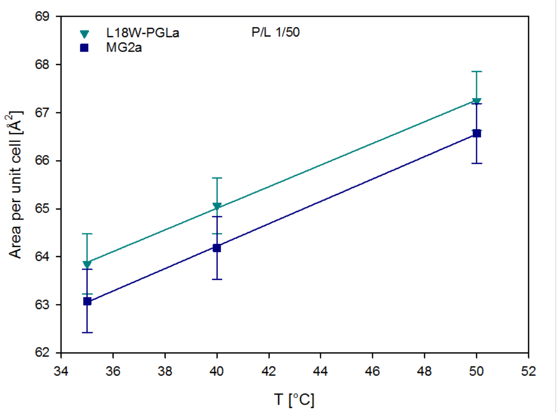


(b)

Figure S11: Temperature dependencies of the area per unit cell for P/L = 1/100 (a) and 1/50 (b). Solid line represent the fits.

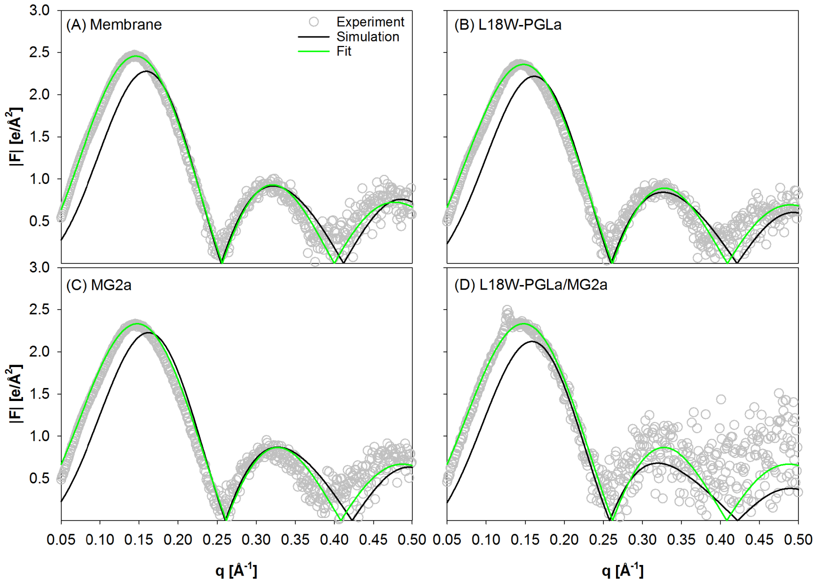


Figure S12: Comparison of experimental form factors for POPE/POPG (3:1 mol/mol) with and without peptides at P/L= 1/200 and 35°C for the pure lipid system (A), L18W-PGLa (B), MG2a (C) and the equimolar mixture (D) to those obtained from MD simulations and global fits. Global fits and simulations are in absolute units while experimental form factors haven been scaled. Note that the global fits include an optimization three additional contrasts from neutron experiments.

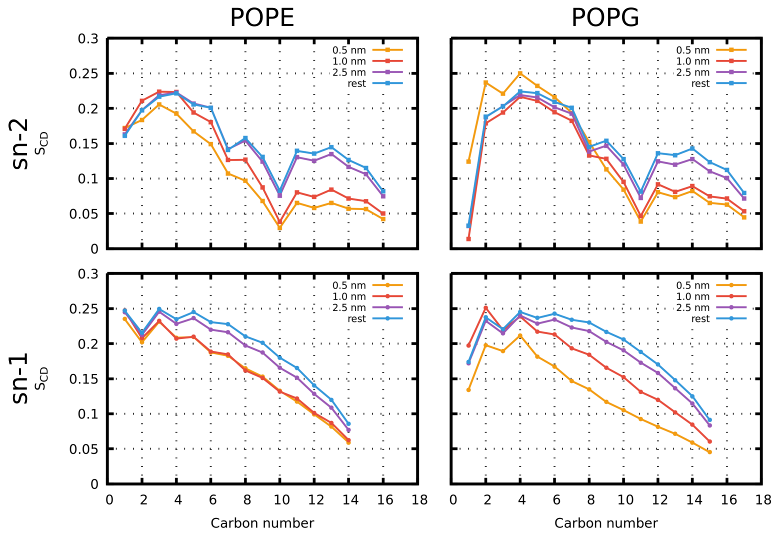


Figure S13: Calculated lipid order parameter of POPE (left) and POPG (right) acyl chains. The data were collected from a 500 ns long all-atom simulation with single MG2a peptide on each membrane leaflet. The distance is given by lipid phosphate to the nearest MG2a peptide C$\alpha$ atom.

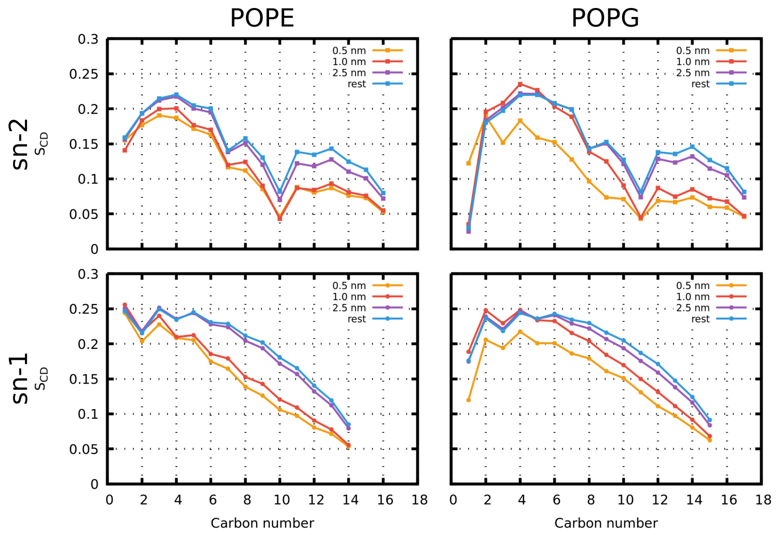


Figure S14: Calculated lipid order parameter of POPE (left) and POPG (right) acyl chains. The data were collected from a 500 ns long all-atom simulation with single L18W-PGLa peptide on each membrane leaflet. The distance is given by lipid phosphate to the nearest L18W-PGLa peptide C$\alpha$ atom.

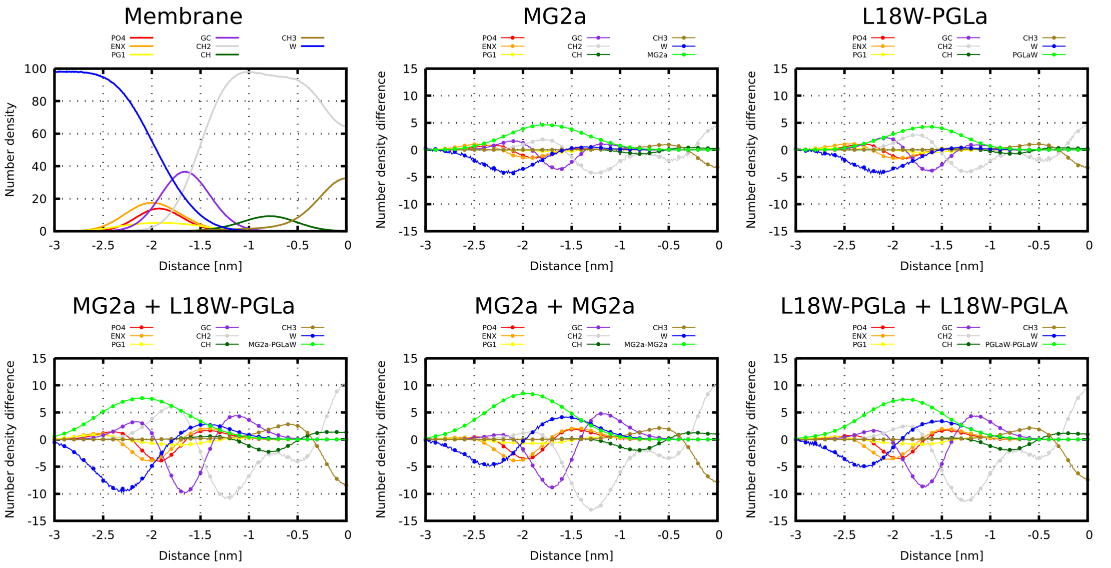


Figure S15: Calculated density profiles from 500 ns long all-atom simulations as a function of distance from the membrane center of mass. (A) Number density of POPE/POPG (3:1 mol/mol) membrane. (B–F) Difference in the number density after the addition of peptides.

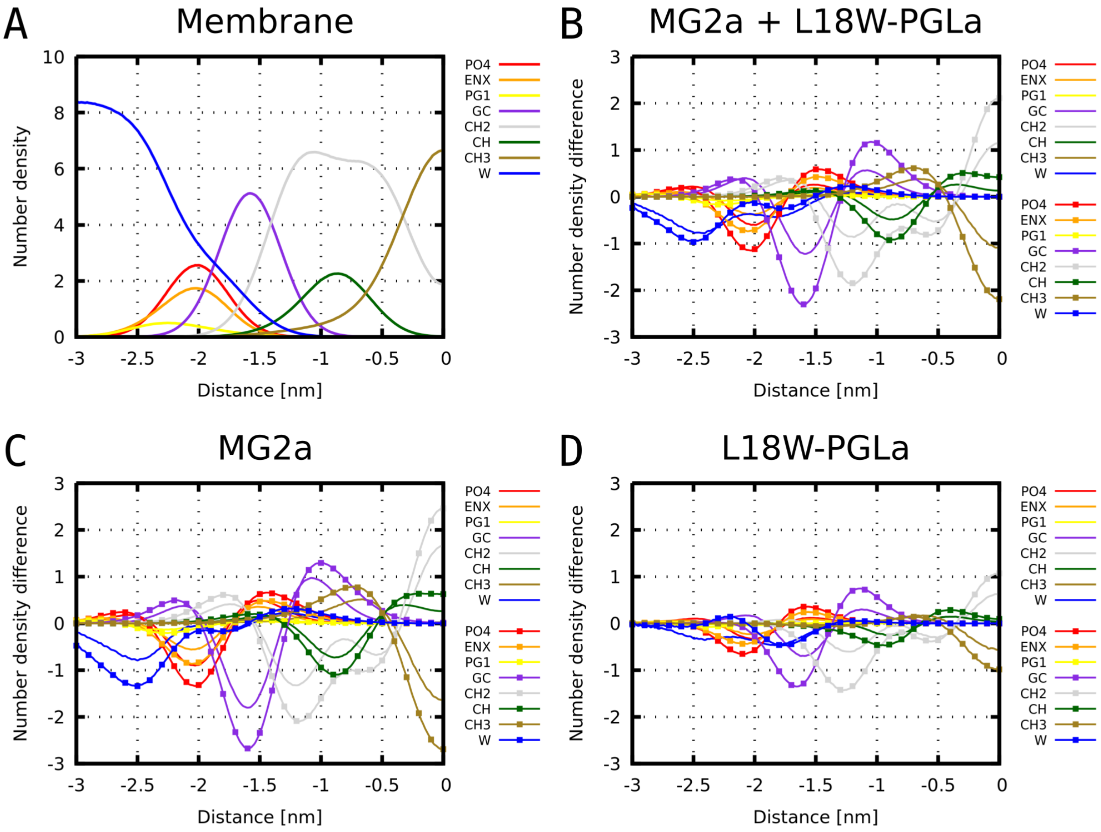


Figure S16: Calculated density profiles from 20 $\mu$s long coarse-grained simulations as a function of distance from the membrane center of mass. (A) Number density of POPE/POPG (3:1 mol/mol) membrane. (B–D) Difference in the number density after the addition of peptides. Systems with P/L = 1/42 and 1/21 are shown with smooth lines and lines with squares, respectively.

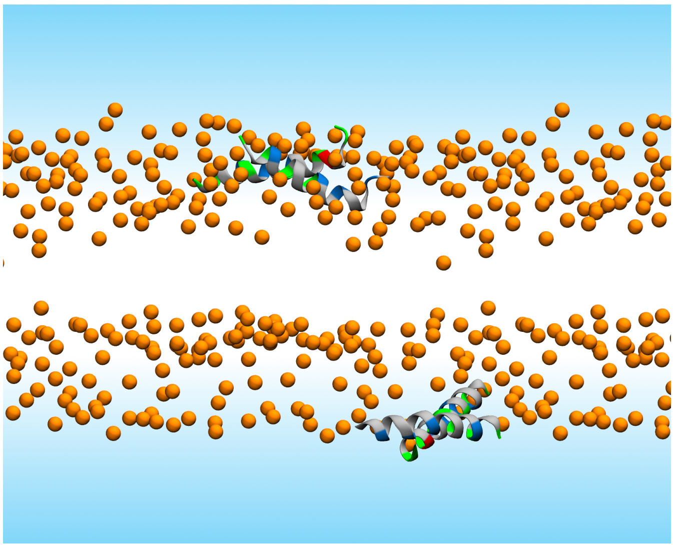

Figure S17: Last snapshots from a 500 ns long MD simulations of MG2a+L18W-PGLa heterodimer adsorbed at the membrane surface. Initially, the peptide pairs were prepared as parallel dimers. Lipid phosphate atoms are shown as orange spheres. Solvent is represented by a blue-shaded area and lipid tails are not shown for clarity. Peptide secondary structure is shown in cartoon representation and colored by residue type. Nonpolar: gray, polar: green, acidic: red, and basic: blue.

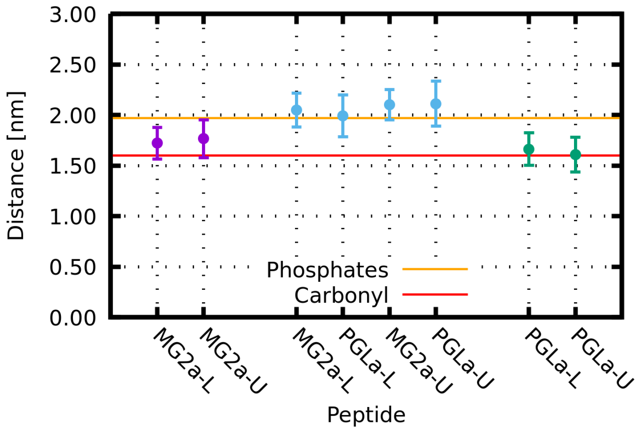


Figure S18: The distance between the peptide and membrane centers of mass. Monomers of (A) MG2a and (C) L18W-PGLa, and heterodimer (B) MG2a+L18W-PGLa are shown. Either (A,C) one monomer or (B) one parallel dimer was adsorbed on (L)ower and (U)pper membrane leaflet. Data are averaged over 500 ns long all-atom trajectory and the bars represent the standard deviation.

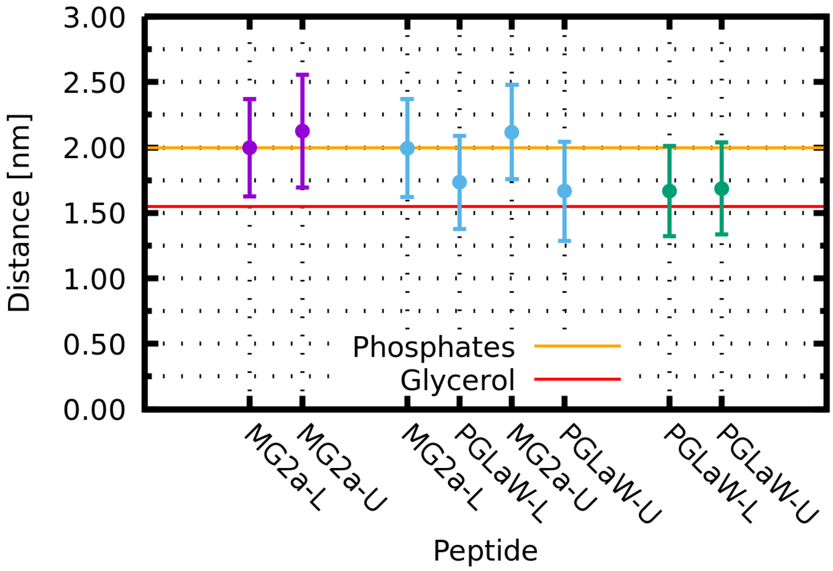


Figure S19: The distance between the peptide and membrane center of mass as obtained from coarse-grained MD simulations. Six peptides (MG2a: purple; 1:1 MG2a+L18W-PGLa: cyan; PGLa: green) were placed on each membrane leaflet (P/L ratio 1/42). Under these conditions, MG2a+L18W-PGLa peptides form parallel heterodimers only occasionally (see Fig. 9 in the main text), causing the L18W-PGLa peptide to be positioned slightly closer towards the membrane surface. The distances are averaged over 20 $\mu$s for all peptides (MG2a or L18W-PGLa) on either the (L)ower or (U)pper membrane leaflets. The bars represent the standard deviation. Orange and red horizontal lines represent the approximate position of phosphate and glycerol groups.

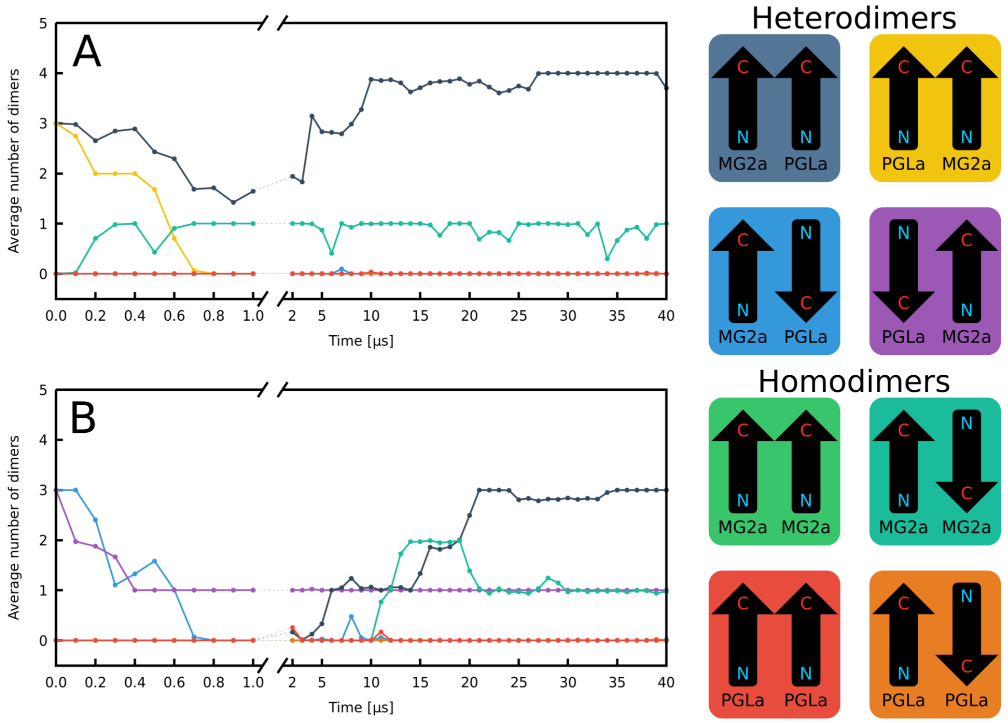


Figure S20: Average number of peptide dimers as a function of time in a system with 12 peptides on each leaflet (P/L = 1/21). Four different heterodimeric starting configurations (each prepared three times) were considered. These configurations are shown in the upper part of the legend. A) On the first leaflet, there were six parallel heterodimers (three of each configuration – gray and yellow) and B) six antiparallel heterodimers (three of each configuration – blue and violet) in the second leaflet. Parallel heterodimers (gray) seem to have the highest stability followed by MG2a homodimers (dark-green). In addition, newly formed dimers were only parallel heterodimers and MG2a antiparallel homodimers, first of which formed more in both A) and B). These results are consistent with simulations starting from random configurations (Fig. 7). Data from the first 1 $\mu$s of the trajectory are averaged every 100 ns. Then, each point is an average from a 1 $\mu$s long trajectory. Color coding of dimers is displayed in the legend on the right side. The instability of parallel PGLa-MG2a heterodimers (yellow) as apposed to MG2a-PGLa heterodimers (grey) is due to the inability to form salt bridge between the residues E19-K15 and stronger tryptophan interactions S23-W18 and I20-W18 of MG2a and L18W-PGLa in such configuration (see Figs. S22 and S23).

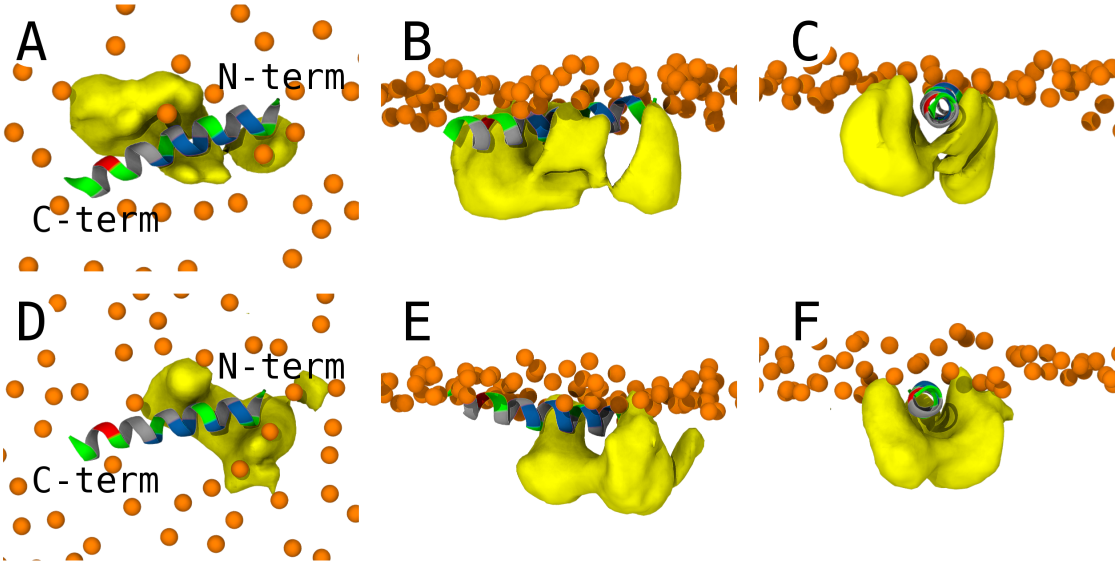


Figure S21: Averaged densities of POPG sn-2 acyl chains from a 500 ns long all-atom MD simulations in the vicinity of MG2a. Lipids with increased order parameter were selected and used for the analysis. In the simulation each membrane leaflet contained one MG2a molecule. Peptides on top (A–C) and bottom (D–E) leaflets were individually analyzed to evaluate the agreement. (A,D) top view on the simulated system, (B,E) side view, and (C,F) view along the peptide long axis. Snapshot color coding: Yellow surfaces represent the average densities of the acyl chain carbons. Lipid phosphate atoms are shown as orange spheres. Solvent and lipid tails are not shown for clarity. The peptide secondary structure is shown in cartoon representation colored by residue type (nonpolar: gray, polar: green, acidic: red, and basic: blue).

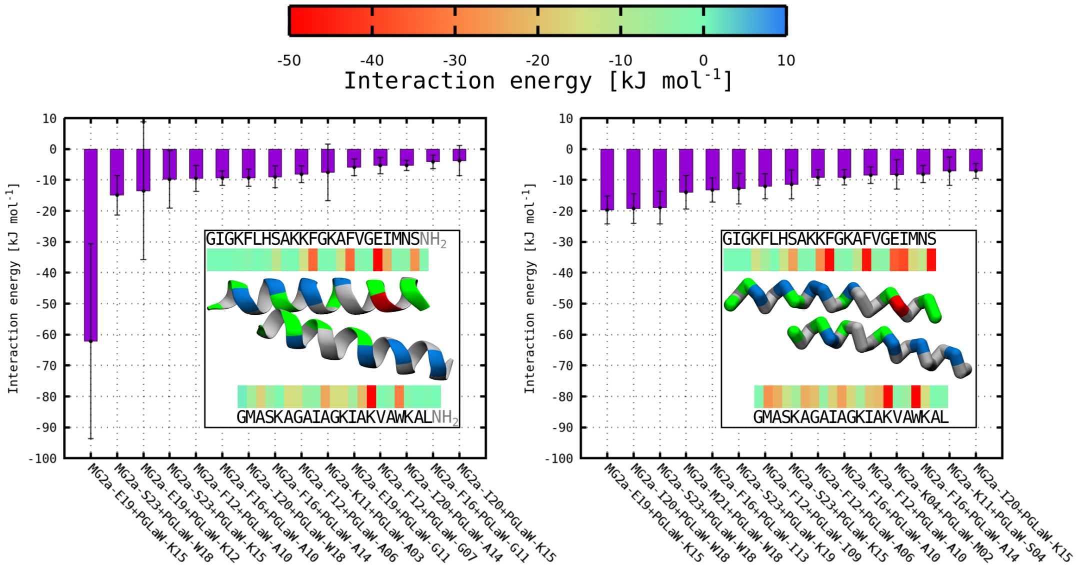


Figure S22: Averaged enthalpic interaction energies between the MG2a and L18W-PGLa heterodimers. Fifteen residue pair interactions with the lowest average energies are shown for both all-atom (left) and coarse-grained (right) simulations. The insets show representative snapshots of MG2a (top) and L18W-PGLa (bottom) peptides in a parallel heterodimer. Color bars show the total average interaction of a residue from one peptide with all residues of the other peptide. The corresponding color scale is presented at the top of the figure. Terminal capping groups were considered as a separate residues according to Amber force field settings (5). Error bars correspond to standard deviations. The large error bars in the all-atom simulations are caused by the possibility of formation of two salt bridges between E19 of MG2a and either K15 and K12 of L18W-PGLa with long residence times. To reach convergence, longer time scale of our all-atom simulation would be required. The peptide secondary structure is shown in cartoon representation colored by residue type (nonpolar: gray, polar: green, acidic: red, and basic: blue).

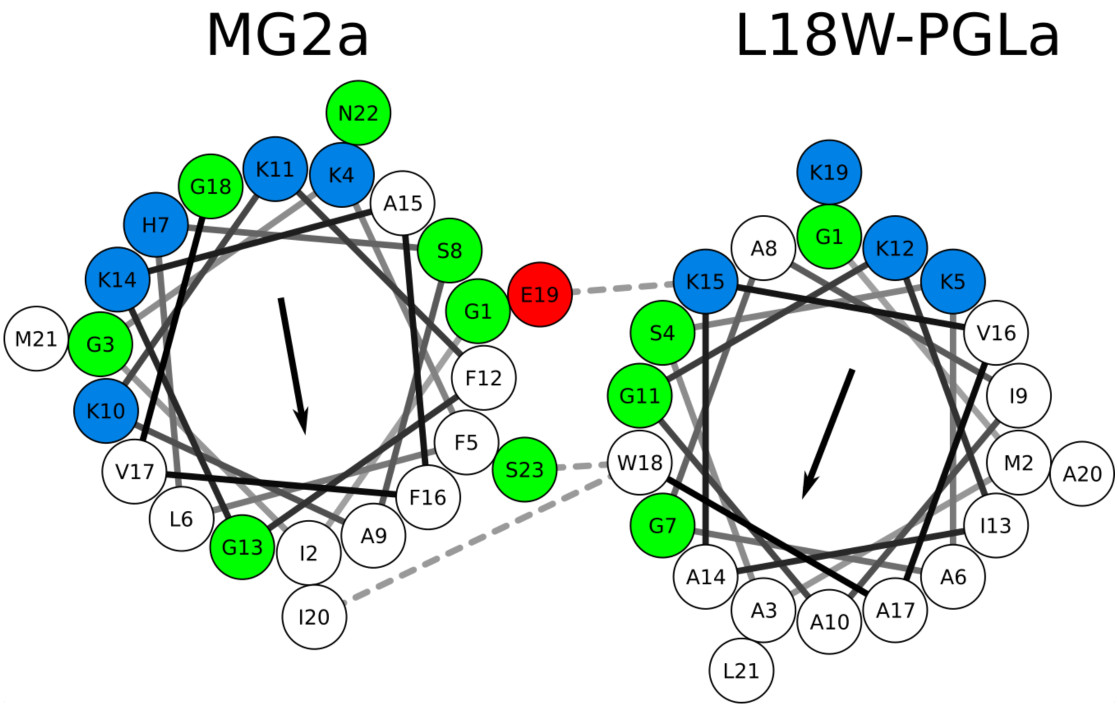


Figure S23: Helical wheels of MG2a and L18W-PGLa arranged in a parallel heterodimer with hydrophobic moments pointing in the same direction. The strongest interactions between peptide residues are marked by dashed lines demonstrating a preference for the displayed conformation. Residues are colored by residue type: nonpolar – white, polar – green, acidic – red, and basic – blue).

### Supplementary Tables

Table S1: Summary of the joint analysis of 100 nm LUVs composed of POPE / POPG (3:1) in the absence and presence of L18W-PGLa, MG2a and their equimolar mixture at P/L = 1/200 and 35°C

| Paramters | POPE/POPG | L18W-PGLa | MG2a | L18W-PGLa / MG2a |
|---|---|---|---|---|
| $z_{CH_2}$ [Å] | 15.34±0.02 | 15.16±0.03 | 14.98±0.02 | 14.68±0.01 |
| $z_{CH}$ [Å] | 9.45±0.22 | 9.53±0.23 | 9.53±0.2 | 9.53±0.21 |
| $z_{GC}$ [Å] | 16.34±0.02 | 16.16±0.03 | 15.98±0.03 | 15.68±0.01 |
| $z_{PO_4}$ [Å] | 20.53±0.02 | 19.96±0.03 | 20.06±0.03 | 20.13±0.03 |
| $z_{ENX}$ [Å] | 22.10±0.02 | 21.53±0.03 | 21.62±0.03 | 21.70±0.03 |
| $z_{PG_2}$ [Å] | 23.45±0.02 | 22.89±0.03 | 22.98±0.03 | 23.05±0.03 |
| $z_P$ [Å] | | 15.76±0.44 | 16.29±0.50 | 15.98±0.43 |
| $\sigma_{CH_3}$ [Å] | 2.64± 0.02 | 2.65±0.02 | 2.65±0.02 | 2.65±0.02 |
| $\sigma_{CH_2}$ [Å] | 3.21±0.05 | 3.19±0.04 | 3.19±0.04 | 3.18±0.04 |
| $\sigma_{CH}$ [Å] | 2.82± 0.04 | 2.82±0.04 | 2.81±0.03 | 2.81±0.04 |
| $\sigma_{GC}$ [Å] | 2.49± 0.03 | 2.48±0.03 | 2.49±0.04 | 2.49±0.03 |
| $\sigma_{PO_4}$ [Å] | 2.47±0.11 | 2.60±0.13 | 2.51±0.16 | 2.72±0.09 |
| $\sigma_{ENX}$ [Å] | 2.81±0.04 | 2.81±0.04 | 2.81±0.04 | 2.80±0.04 |
| $\sigma_{PG_2}$ [Å] | 3.15±0.04 | 3.16±0.04 | 3.16±0.04 | 3.16±0.04 |
| $\sigma_P$ [Å] | | 4.02±0.14 | 4.04±0.15 | 4.00±0.14 |
